# Adolescent alcohol consumption differentially alters sexually-divergent behaviors and infralimbic cortex function in mice

**DOI:** 10.1101/2025.06.05.658112

**Authors:** Laurel R. Seemiller, Hayreddin Said Ünsal, Yiyan Xie, Md Shakhawat Hossain, Avery R. Sicher, Keith R. Griffith, Kaleah Tuttle, Bryce Johnson-Dendy, Emily P. McDonald, Lauren N. Beloate, Patrick J. Drew, Nanyin Zhang, Nicole A. Crowley

## Abstract

The cortex is one of the last brain regions to fully mature, making it particularly sensitive to early life drug use. Both human and rodent studies find long-lasting behavioral changes after adolescent alcohol exposure that implicate underlying disruptions in cortical development, including structural abnormalities and altered functional connectivity. Few rodent studies have been conducted across biological scales (from single cell, whole brain network, to behavior) to understand the comprehensive implications of these disruptions. We assessed how adolescent binge-like alcohol consumption in a drinking in the dark (DID) model affected adult alcohol consumption, behavioral exploration, brain-wide functional connectivity, and intrinsic cellular excitability in mice. Only female mice exposed to alcohol during adolescence exhibited aversion-resistant alcohol preference, as well as increased exploration in the elevated plus maze. Resting state neuroimaging revealed broad sex-specific changes in brain-wide networks. Finally, we identify *ex vivo* intrinsic excitability alterations in key brain regions known to be susceptible to adolescent perturbations identified in neural networks, with greater effects in males, allowing biological and mechanistic targets to be further explored for future study and interventions. Our results show that females may be more vulnerable to lasting addiction-related behavioral and brain connectivity consequences of adolescent alcohol exposure, corroborating human and rodent studies demonstrating that adolescent alcohol use may increase risk for adult alcohol use more strongly in females. We further show that while males do not show identical behavioral changes, they critically show alterations in intrinsic excitability and cellular function, suggesting divergent sex-specific adaptations to adolescent binge drinking.

## 1. INTRODUCTION

Adolescent alcohol use is associated with increased adult alcohol use as well as an increased risk of developing an alcohol-related disorder [1–3]. This risk persists throughout the lifetime and is stronger in females than in males [3]. Consequences of adolescent alcohol exposure are thought to be driven by disturbances in the natural development of both morphological characteristics and functional wiring properties of the brain. While normative adolescent brain development requires selective pruning of neuronal connections and cortical thinning [4], adolescent alcohol use can disrupt these key developmental processes [5]. Complementary lines of evidence show disruptions in functional connectivity between the default-mode network and an intrinsic functional network implicated in regulating emotional responses [6] as well as within cortical and striatal networks [7], suggesting potential network mechanisms for persistent behavioral consequences of adolescent alcohol use. Vulnerability to development of an alcohol use disorder (AUD) can also be exacerbated by other psychiatric conditions, such as anxiety [8] (which is heightened after alcohol use [9]) suggesting reciprocal behavioral and physiological feedback between substance use and mood disorders [10].

Despite known consequences of adolescent binge drinking and recent novel insights into alcohol-driven brain connectivity changes, further work is needed to bridge the gap between human epidemiological studies and mechanistic biological brain changes observed in rodent studies. Here, we aimed to narrow this divide by examining the long-term consequences of voluntary alcohol consumption during adolescence in a mouse model. Male and female C57BL/6J mice received intermittent access to alcohol in a Drinking in the Dark (DID) paradigm [11] throughout adolescence. After DID, they were tested for changes in aversion-resistant alcohol preference, exploratory and anxiety-like behaviors, whole-brain resting state functional connectivity [12], and *ex vivo* intrinsic excitability of selective cortical and subcortical excitatory neuronal populations in adulthood. We found binge drinking-induced changes in alcohol preference and exploratory and anxiety-like behavior, altered brain-wide and region-specific functional connectivity, and changes in region-specific pyramidal cell excitability, all of which showed marked sex differences. Our data suggest alterations in cortical networks induced by voluntary alcohol consumption may underlie adulthood shifts in risk-taking and executive-decision making behaviors in a well-validated mouse binge drinking model, consistent with what has been proposed in the human literature [13]. Our current work further bridges the gap between human epidemiological and behavioral studies and preclinical mechanistic biological studies via a shared non-invasive imaging dataset and allows for synthesis across human and animal studies.

## 2. MATERIALS AND METHODS

### 2.1 Animals

Male and female C57BL/6J (JAX #000664) mice were bred in-house for experiments beginning during adolescence (Bar Harbor, ME, USA; see **Supplemental Materials** for adult experiments). On postnatal day (PND) 21, mice were weaned and single-housed on a reversed light cycle (lights off at 7:00 AM). They were habituated to these conditions for at least one week prior to experiments. Mice had *ad libitum* access to food (LabDiet 5053, Lab Diet, St. Louis, MO, USA), and *ad libitum* access to water outside of DID testing. All procedures were conducted in accordance with the NIH Guide for the Care and Use of Laboratory Animals and were approved by the Pennsylvania State University Institutional Animal Care and Use Committee.

### 2.2 Drinking in the Dark (DID)

DID was performed as described previously [11,14–16]. On PND 29 (+/-1 day) beginning 3 hr into the dark cycle, mice had access to 20% v/v ethanol (EtOH; Koptec, Decon Labs, King of Prussia, PA) in H_2_O (tap water) instead of their normal H_2_O bottle. Alcohol access lasted 2 hr for 3 days and 4 hr for the fourth day. After 3 days of abstinence, the DID cycle was repeated 4 times, with the last binge day on PND 53 (+/-1 day; see **Supplemental Materials** for ages during adulthood behavior experiments). Control subjects received access to H_2_O. Mice were weighed once weekly.

### 2.3 Behavioral testing

Adulthood EtOH preference was measured as published previously [16,17]. Mice were habituated to H_2_O tubes for one week before receiving continuous access to H_2_O and EtOH in a two-bottle choice (2BC) paradigm. After preference for EtOH was established, EtOH was adulterated with increasing concentrations of quinine (see **Supplemental Materials** for more details).

In a separate cohort of mice dedicated to behavioral testing, behaviors in the open field test (OFT) and elevated plus maze (EPM) were assessed roughly 30 days (or 24 hr, see **Supplemental Materials**) after DID. Mice habituated to testing rooms for at least one hour prior to a 5 min OFT test. At least one hour after the OFT test, mice underwent a subsequent 5 min EPM test. Behavior was video-recorded and analyzed using DeepLabCut [18] and MATLAB (see **Supplemental Materials**).

### 2.4 Functional magnetic resonance imaging (fMRI)

Beginning four days prior to imaging, mice were briefly anesthetized with 3% isoflurane, secured in a restrainer, allowed to awake from anesthesia, and restrained for 15, 30, 45, and 60 min on acclimation days 1-4, respectively, while fMRI noises were played. Mice were anesthetized again before being removed from the restrainer. On PND 83-88 (at least 30 days after DID), awake resting state imaging was performed using a 7T MRI system and Bruker Console (Billerica, MA) using Paravision 7.0.0. Each scan lasted 10 min and collected 400 volumes, with three scans per mouse. Framewise displacement (FD) was calculated to quantify motion [19]: FD = ∣ Δx ∣ + ∣ Δy ∣ + ∣ Δz ∣ + r * (∣ Δα ∣ + ∣ Δβ ∣ + ∣ Δγ ∣). Motion calculation and correction were performed using the Statistical Parametric Mapping (SPM12) package (http://www.fil.ion.ucl.ac.uk/spm/). A total of 62 regions of interest (ROIs) were defined based on the Allen Mouse Brain Atlas [20]. An ROI-based approach specifically tested Pearson correlations between average activity time course between each ROI pair. Connectivity matrices were calculated using Fisher’s z-transformed Pearson’s correlation coefficients within each mouse. Methodological details for fMRI can be found in **Supplemental Materials**.

### 2.5 *Ex vivo* patch clamp electrophysiology

Prior to electrophysiology, all mice underwent EPM (30 days after DID and 1-6 days prior to sacrifice) as described above (EPM data not included). Mice were transferred from the vivarium to the laboratory and habituated undisturbed in a fume hood for 1 hr prior to slicing (see **Supplemental Materials** for additional methods). Whole-cell patch-clamp electrophysiology was performed as previously described [15,17,21,22] to assess passive properties and intrinsic excitability of pyramidal cells in multiple brain regions. A maximum of 3 cells per region were recorded from each mouse. Signals were digitized at 10 kHz and filtered at 4 kHz with a MultiClamp 700B amplifier. Recordings were analyzed using Clampfit 11.3 software (Molecular Devices, Sunnyvale, CA, United States).

### 2.6 Statistics

*Behavior and electrophysiology analysis:* Statistical analysis was performed in GraphPad Prism (version 10; La Jolla, CA, USA) and MATLAB (R2021a; The MathWorks Inc., Natick, MA, United States) and figures were made in GraphPad Prism. A significance threshold of α = 0.05 was used for all analyses. Two– and three-way ANOVA were used, employing a mixed model for repeated measures. Sex and alcohol interactions were followed up with post-hoc Fisher’s LSD or t-tests with a Bonferroni correction between alcohol and control groups within each sex. Sexes were analyzed separately only for 2BC experiments due to differences in adolescent alcohol consumption (see **Supplemental Methods**). Additional generalized linear mixed-effects (GLME) modeling in alcohol-exposed mice explored the relationship between total alcohol consumption and adult outcomes using the fitglme toolbox in MATLAB and accounted for litter and behavioral cohort as random effects: [outcome variable] ∼ 1 + Sex*EtOH + (1|Litter) + (1|Cohort). The link function was identity or log for variables with normal and Poisson distributions, respectively. When sex and alcohol interactions were detected, they were followed up with post-hoc additional GLME models within each sex: [outcome variable] ∼ 1 + EtOH + (1|Litter) + (1|Cohort). More statistical details can be found in **Supplemental Materials**.

*fMRI analysis:* Statistical analysis was performed and figures were made in MATLAB version R2023b (The MathWorks Inc., Natick, MA, United States). Runs with over 10% of volumes discarded due to motion artifacts were excluded from final analysis. The effects of adolescent alcohol consumption, sex, and their interaction on functional connectivity were assessed using a linear mixed model, with subject as a random effect. Averaged seed-to-whole brain connectivity is quantified by averaging the functional connectivity between the seed ROI and all other ROIs.

## 2 RESULTS

### 3.1 Adult preference for alcohol and quinine-adulterated alcohol after adolescent drinking

Adolescent male and female mice received intermittent access to alcohol in a DID paradigm (alcohol consumption and body weights described in **Supp. Fig. 1**). Beginning approximately 30 days after adolescent drinking, adult mice received continuous access to a bottle of alcohol containing increasing concentrations of quinine alongside a bottle of water (timeline in **Fig 1A**). Adult females that consumed alcohol during adolescence exhibited stronger preference for alcohol and quinine-adulterated alcohol than females that only had water during adolescence (F_1,18_ = 4.7, *p* = 0.0441; **Fig 1B**). Adult female alcohol, water, total fluid consumption, and adult body weight were not significantly altered by adolescent alcohol exposure (**Fig 1C-D, Supp. Fig. 2**). In adult males, adolescent alcohol consumption did not significantly influence adult alcohol preference, alcohol consumption, water consumption, total fluid consumption, or body weight (**Fig 1E-G, Supp. Fig. 2**). Total adult alcohol consumed prior to the addition of quinine was positively associated with total alcohol consumed with quinine (**Supp. Fig. 2**). Overall, adolescent alcohol consumption increased adult preference for alcohol and quinine-adulterated alcohol in females but not in males.

**Figure 1:**
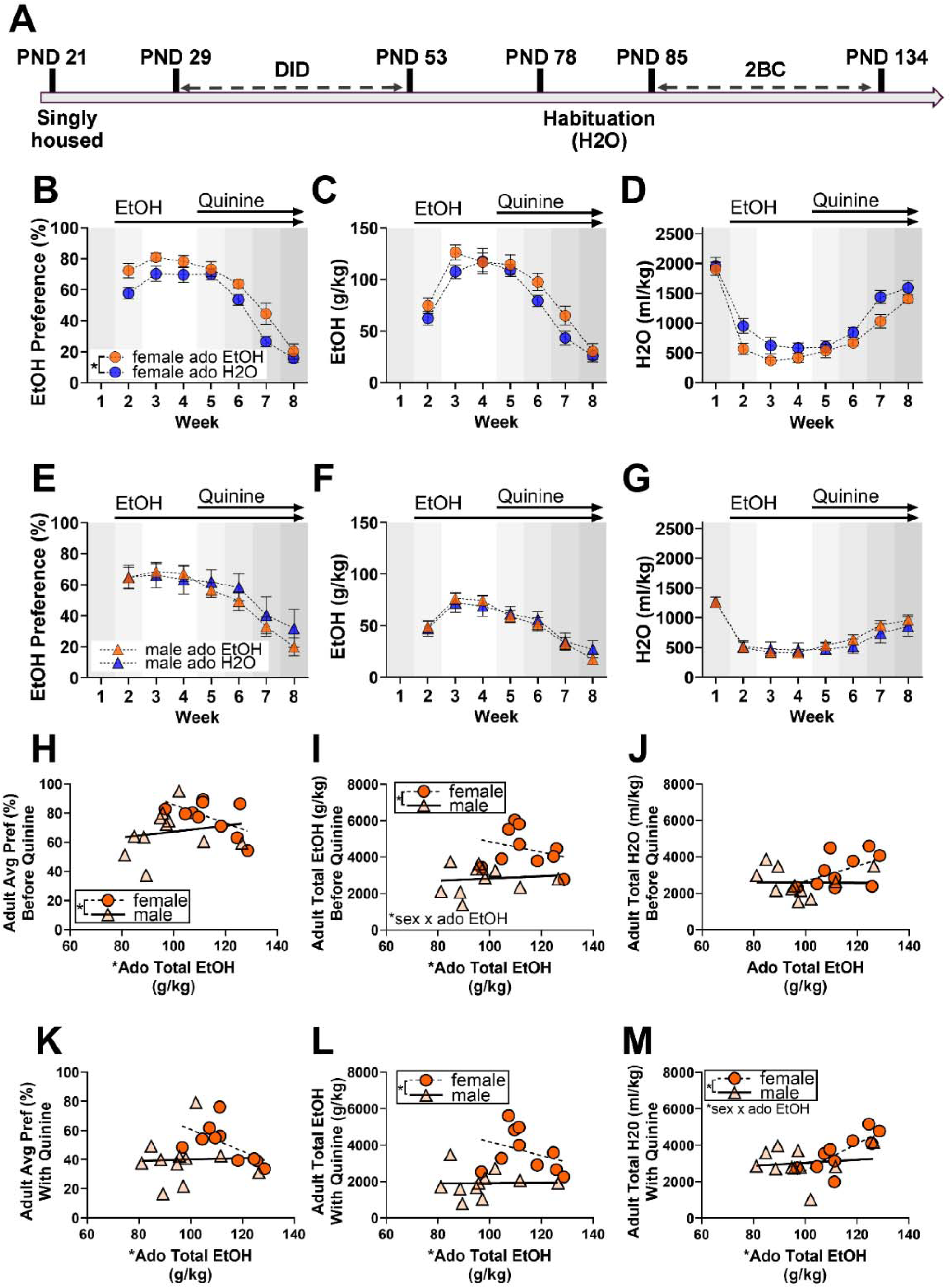
Adolescent alcohol consumption increased adult preference for alcohol and quinineadulterated alcohol in females and not in males. (**A**) Experimental timeline for drinking in the dark (DID) effects on adult aversion-resistant alcohol preference via two-bottle choice (2BC). **(B)** EtOH preference in females. **(C)** EtOH g/kg consumed by females. **(D)** Water consumed in ml/kg by females. **(E)** EtOH preference in males. **(F)** EtOH g/kg consumed by males. **(G)** Water consumed in ml/kg by males. **(H)** Average EtOH preference compared to adolescent EtOH consumption in females. **(I)** Total EtOH consumption during 2BC in g/kg compared to adolescent EtOH consumption by females. **(J)** Total water consumption during 2BC in ml/kg compared to adolescent EtOH consumption in females. **(K)** Average EtOH preference with quinine compared to adolescent EtOH consumption in males. **(L)** Total adult EtOH consumption with quinine during 2BC in g/kg compared to adolescent EtOH consumption in males. **(M)** Total water consumption alongside quinine testing during 2BC in ml/kg compared to adolescent EtOH consumption in males. Ado = Adolescent. Week 1 represents consumption of H_2_O only, Week 2 includes 3-7% EtOH, Week 3-4 represents 10% EtOH, and Weeks 5-8 represent 10% EtOH with increasing concentrations of quinine (0.03, 0.1, 0.3, and 1mM). Plots labeled as “Before Quinine” represent data from Weeks 1-4, and plots labeled as “With Quinine” represent data from Weeks 5-8. In scatter plots, a simple linear regression was used to generate lines of best fit and aid interpretation but were independent from statistical analysis. Data are shown as mean +/− SEM. * *p* <0.05. n=9-11/sex/treatment.

**Figure 2:**
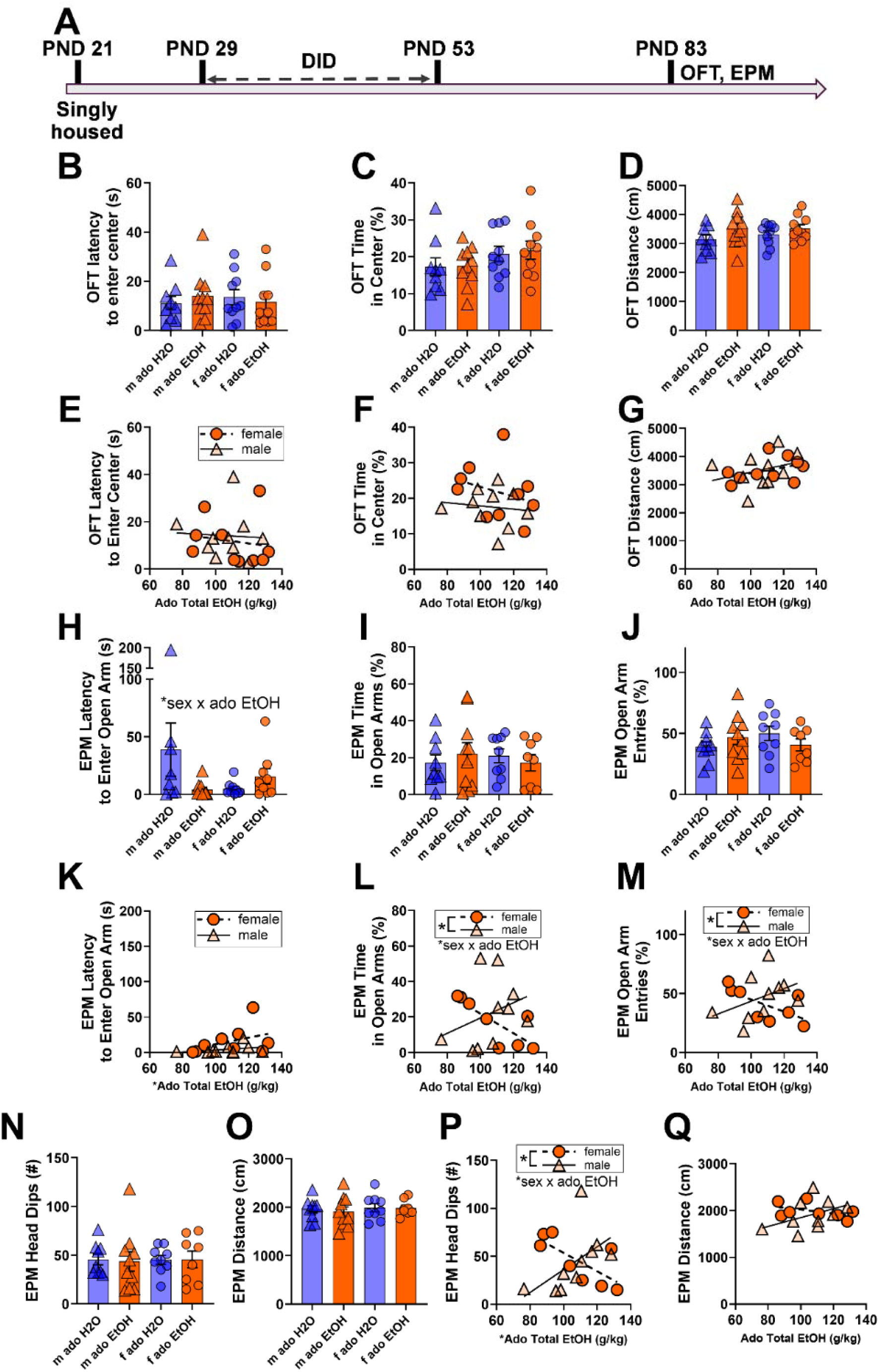
Adolescent alcohol consumption had sex-specific effects on adult exploratory behaviors. (**A**) Experimental timeline. **(B)** OFT latency to enter center of arena. **(C)** OFT percent time spent in center of arena. **(D)** OFT total distance traveled. **(E)** OFT latency to enter center compared to adolescent EtOH consumption. **(F)** OFT percent time in center compared to adolescent EtOH consumption. **(G)** OFT total distance traveled compared to adolescent EtOH consumption. **(H)** EPM latency to enter an open arm. **(I)** EPM percent time spent in open arms. **(J)** EPM percent open arm entries. **(K)** EPM latency to enter an open arm compared to adolescent EtOH consumption. **(L)** EPM percent time spent in open arms compared to adolescent EtOH consumption. **(M)** EPM percent open arm entries compared to adolescent EtOH consumption. **(N)** EPM number of head dips. **(O)** EPM total distance traveled. **(P)** EPM number of head dips compared to adolescent EtOH consumption. **(Q)** EPM total distance traveled compared to adolescent EtOH consumption. M = male; f = female; ado = adolescent. In scatter plots, a simple linear regression was used to generate lines of best fit and aid interpretation but were independent from statistical analysis. Data are shown as mean +/− SEM. *p<0.05. n=8-10/sex/treatment.

To directly compare sexes while accounting for differences in adolescent alcohol consumption, we used a GLME model. Before quinine testing, total adolescent alcohol consumption was negatively associated with adult average alcohol preference (ado EtOH: –0.0092989 ± 0.0040242, p = 0.0337) and consumption (ado EtOH: –77.03 ± 19.392, p = 0.0010). Males exhibited lower adult alcohol preference (quinine male: –1.22 ± 0.51557, p = 0.0301; **Fig 1H**) and consumption (male: –7300.1 ± 2334.3, p = 0.0061; **Fig 1I**) than females. Total adult alcohol consumption without quinine was also influenced by an interaction of sex and adolescent alcohol (male x ado EtOH: 46.756 ± 21.202, p = 0.0415). Post-hoc analyses within sexes confirmed an effect of adolescent alcohol in males (ado EtOH: –20.486 ± 8.4987, p = 0.0392) but not in females. Adult total water consumption prior to quinine testing was similar across groups (**Fig 1J**). After quinine was included (weeks 5–8), total adolescent alcohol consumption was negatively associated with adult average preference (ado EtOH: – 0.010501 ± 0.0043085, p = 0.0261; **Fig 1K**) and consumption (ado EtOH: –85.924 ± 22.866, p = 0.0016) of alcohol with quinine. Sex differences were not observed for preference, but adult males consumed less alcohol with quinine than adult females (male: –7122.3 ± 2759.8, p = 0.0194; **Fig 1L**). Adolescent alcohol consumption was positively associated with adult water consumption (ado EtOH: 73.618 ± 23.207, p = 0.0056), adult males consumed more water than adult females (male: 6981.3 ± 3163.4, p = 0.0414), and sex and adolescent alcohol consumption interacted to influence adult water consumption (male x ado EtOH: –65.655 ± 29.139, p = 0.0377; **Fig 1M**). Post-hoc analyses found a significant positive effect of adolescent alcohol consumption on adult water consumption during quinine testing in females (ado EtOH: 73.618 ± 19.258, p = 0.0051) but not in males. Additionally, total fluid consumption and average adult body weight were not impacted by sex or total adolescent alcohol consumption (**Supp. Fig. 2**). To summarize, total adolescent alcohol consumption had a negative association with average adult preference and alcohol consumption before and during the quinine test.

### 3.2 Adult exploration in the OFT and EPM after adolescent drinking

In a separate cohort of mice, we tested adult exploratory behaviors after adolescent alcohol consumption (**Fig 2A**). OFT measures were not affected by sex or adolescent alcohol exposure (**Fig 2B-G**). However, in the EPM, adolescent alcohol consumption and sex significantly interacted to alter latency to enter an open arm (F_1,32_ = 4.4, p = 0.0446; **Fig 2H**), though post-hoc tests within sexes did not reveal differences across treatment groups. Time spent in open arms (**Fig 2I**) and open arm entries (**Fig 2J**) were not different across alcohol or sex groups. Additional analyses tested the association between total adolescent alcohol consumption and adult behaviors in only alcohol-exposed mice. No effects were seen for latency to enter the open arm of the EPM (**Fig 2K**), but this revealed an interaction of adolescent alcohol consumption and sex on time spent in the open arm (male x ado EtOH: 0.9827 ± 0.4244, p = 0.0363; **Fig 2L**) and open arm entries (male x ado EtOH: 1.0538 ± 0.4220, p = 0.0256; **Fig 2M**). Post-hoc analyses within sexes revealed reduced time spent in the open arm (ado EtOH: –0.6544 ± 0.0290, p < 0.0001) and open arm entries (ado EtOH: – 0.5535 ± 0.1947, p = 0.0295) with increasing adolescent alcohol consumption levels in females and not in males. Further, total adolescent alcohol consumption was associated with latency to enter the open arm (ado EtOH: 0.0463 ± 0.01146, p = 0.0011). Sex was associated with time spent in open arms (male: –100.69 ± 45.956, p = 0.0459) and entries into open arms (male: –106.73 ± 45.694, p = 0.0349). Further, head dips (**Fig 2N**) and distance traveled (**Fig 2O**) were not affected by treatment or sex. However, a significant interaction of total adolescent alcohol consumption and sex was detected for head dips (male x ado EtOH: 2.1466 ± 0.6536, p = 0.0054) in addition to a significant influence of total adolescent alcohol exposure (ado EtOH: –0.9572 ± 0.4410, p = 0.0477) and sex (male: –232.09 ± 70.78, p = 0.0055; **Fig 2P**). Post-hoc analyses within sexes showed significant reductions in head dips due to total alcohol exposure within females, and not males (ado EtOH: –1.0553 ± 0.2346, p = 0.0041). No effects of sex or total alcohol consumed were seen for distance traveled in the EPM (**Fig 2Q**). In summary, across multiple measures, total adolescent alcohol consumption decreased EPM exploratory behaviors in females and not in males.

To understand whether the effects of alcohol were specific to adolescents or the 30-day post-DID timepoint, we investigated how the same DID paradigm affected adolescents 24 hr post-DID (**Supp. Fig. 3**) and adults 30 days post-DID (**Supp. Fig. 4**). There were only minor effects of alcohol on behavior in these experiments, suggesting that the adolescent period is uniquely vulnerable to alcohol and some behavioral effects of adolescent alcohol exposure may take time to emerge.

**Figure 3.**
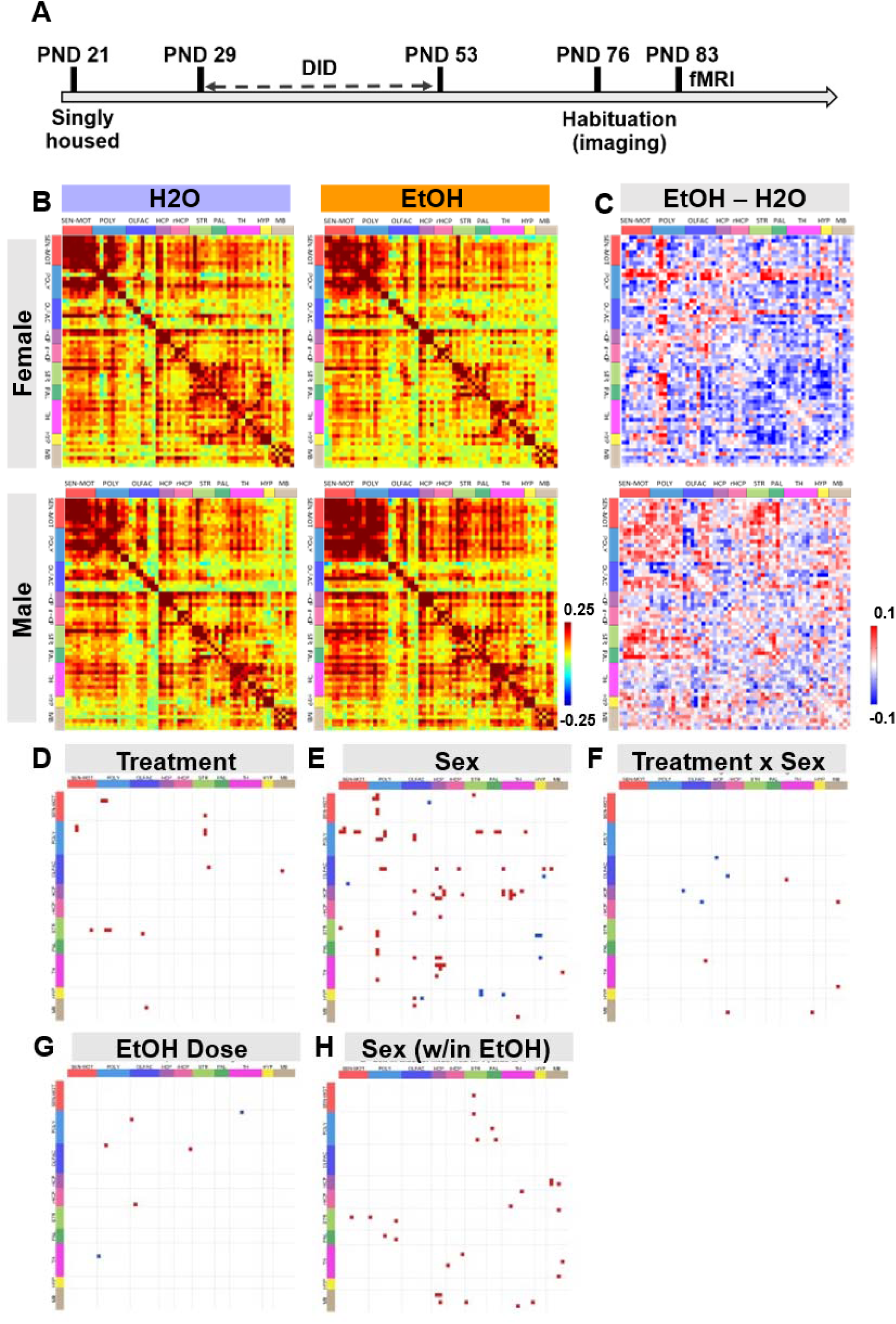
Whole brain functional connectivity differs across sex and due to adolescent alcohol consumption. (**A**) Experimental timeline. **(B)** Average resting state functional connectivity (RSFC) matrices for sex and alcohol groups. The color bar (± 0.25 range) represents the z-score values for the corresponding RSFC. **(C)** Difference of RSFC between alcohol and control groups for each sex. The red-blue color bar (± 0.1 range) shows the z-score differences for the increased or decreased average RSFC in adolescent alcohol-exposed mice in comparison to the controls. **(D)** Significant differences in RSFC between treatment groups. Red indicates greater RSFC in alcohol-treated mice. **(E)** Significant differences in RSFC between sexes. Red indicates greater RSFC in males, and blue indicates greater RSFC in females. **(F)** Significant differences in RSFC due to an interaction of alcohol and sex. Red indicates greater effects of alcohol in males, and blue indicates greater effects of alcohol in females. **(G)** Significant effects of total alcohol consumption within alcohol-exposed mice. Red indicates positive effects of total alcohol on RSFC, and blue indicates negative effects of total alcohol on RSFC. **(E)** Significant effects of sex on RSFC within alcohol-exposed mice. Red indicates greater RSFC in alcoholexposed males. An adjusted significance threshold of p<0.005 was used for pairwise RSFC comparisons. n=11 mice/sex/treatment.

**Figure 4.**
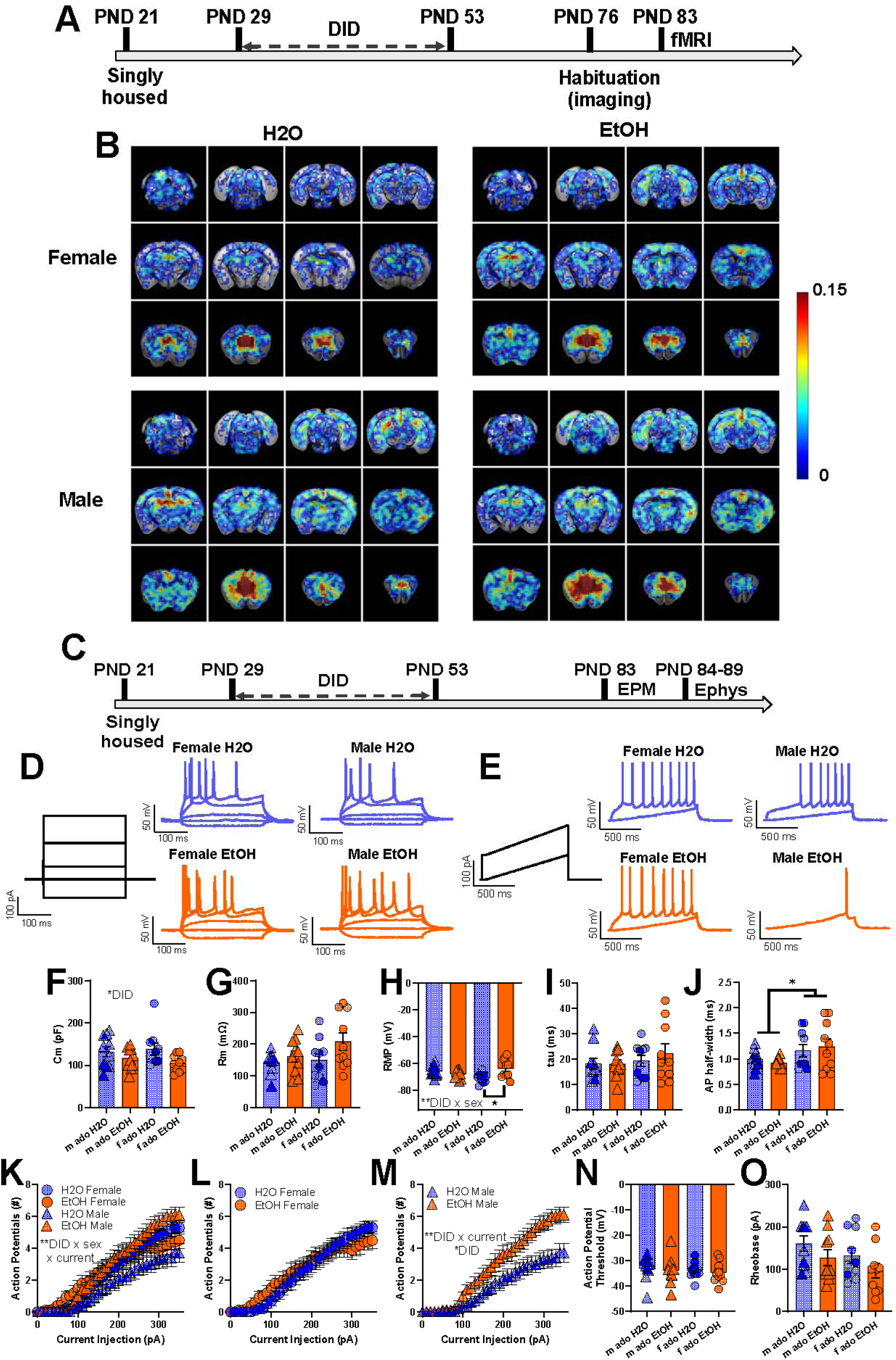
Adolescent alcohol consumption reduces infralimbic cortex functional connectivity and pyramidal cell excitability. (**A**) Experimental timeline for fMRI experiments. **(B)** Infralimbic seedbased functional connectivity maps. The color bar (0.15 range) represents average RSFC correlation values between the infralimbic and all other voxels. **(C)** Experimental timeline for patch clamp electrophysiology experiments testing the intrinsic excitability of infralimbic pyramidal cells. **(D)** Representative traces from current-induced firing experiments. **(E)** Representative traces from rheobase recordings. **(F-J)** Cellular capacitance, membrane resistance, resting membrane potential (RMP), time constant (tau), and action potential (AP) half-width**. (K-M)** Current-induced firing of cells at RMP, with sexes visualized together **(K)** and separately **(L-M)**. **(N-O)** Action potential threshold and rheobase of cells at RMP. M = male; f = female; ado = adolescent. Data are shown as mean +/− SEM. *p<0.05. **p<0.01. n=11 mice/sex/treatment for fMRI and n=7-9 mice/sex/treatment for electrophysiology.

### 3.3 Adolescent binge drinking causes sex-specific alterations in adult whole-brain functional connectivity

An additional cohort of mice underwent resting state functional magnetic resonance imaging (rs-fMRI) roughly 30 days after adolescent drinking (**Fig 3A**). The mouse brain was partitioned to 62 bilateral regions of interest (ROIs) anatomically defined in the Allen Mouse Brain Atlas [20] (see **Supp. Fig. 5** for ROI descriptions). Globally, females showed a strong brain-wide decrease in overall functional connectivity post-binge drinking (**Fig 3B-C**), while this pattern was reversed in males (**Fig 3B-C**). Alcohol consumption, sex, and interactions of alcohol and sex significantly altered resting state functional connectivity in region-specific manners (**Fig 3D-H**). First, region-specific analyses were focused on the prefrontal cortex (PFC) due to its known plasticity during adolescence. Of all subregions of the medial PFC, only the infralimbic cortex showed altered RSFC due to adolescent alcohol exposure, specifically in female mice (**Supp. Fig. 6**). Together, functional imaging data demonstrates a robust sex-dependent and dose-dependent alteration in whole brain resting state functional connectivity, consistent with sex differences in behavioral profiles.

**Figure 5.**
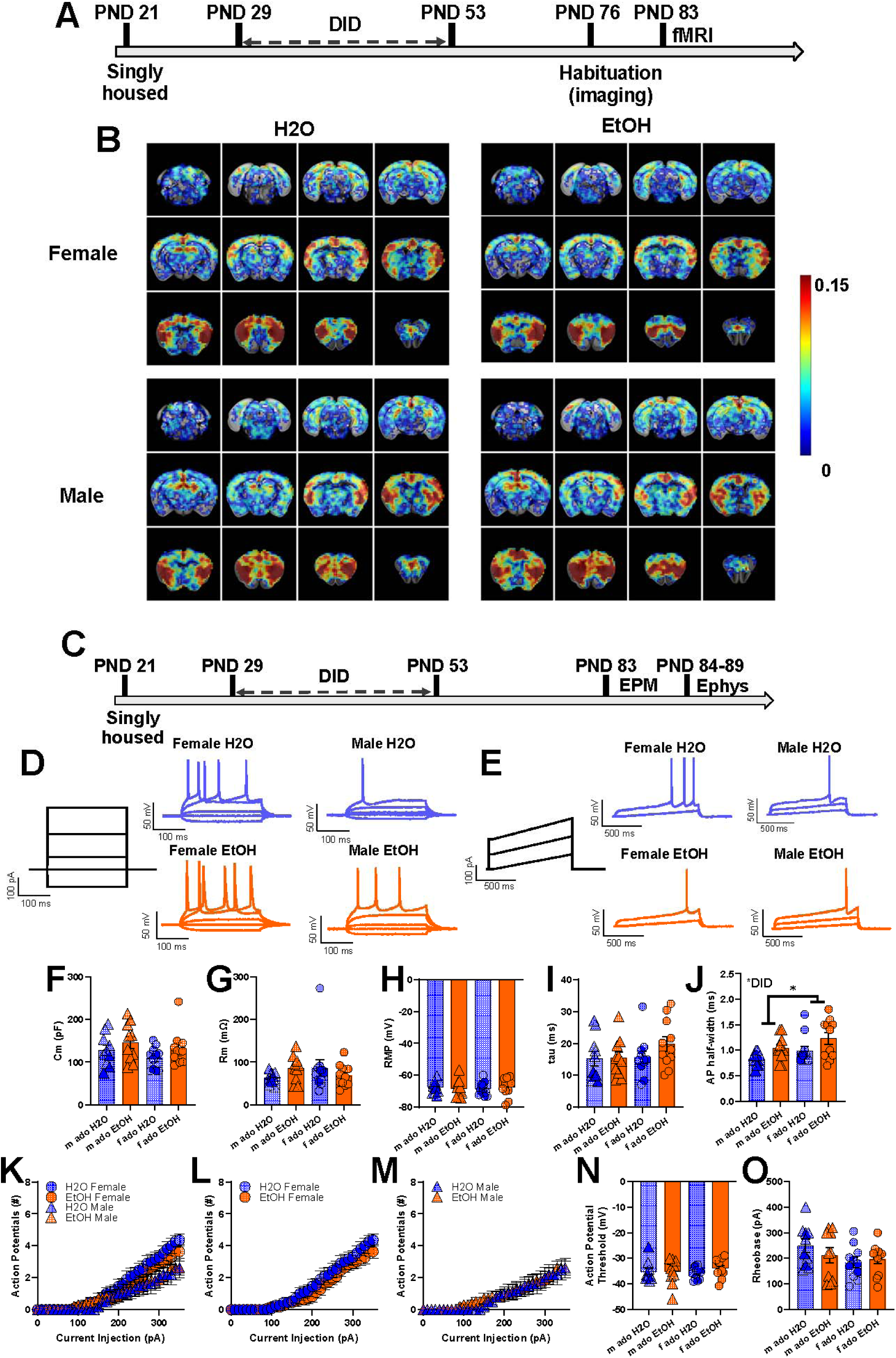
Adolescent alcohol consumption alters anterior insular cortex functional connectivity and pyramidal cell excitability. (**A**) Experimental timeline for fMRI experiments. **(B)** Anterior insular cortex seed-based functional connectivity maps. The color bar (0.15 range) represents average RSFC correlation values between the anterior insular cortex and all other voxels. **(C)** Experimental timeline for patch clamp electrophysiology experiments testing the intrinsic excitability of anterior insular cortex pyramidal cells. **(D)** Representative traces from current-induced firing experiments. **(E)** Representative traces from rheobase recordings. **(F-J)** Cellular capacitance, membrane resistance, resting membrane potential (RMP), time constant (tau), and action potential (AP) half-width**. (K-M)** Current-induced firing of cells at RMP, with sexes visualized together **(K)** and separately **(L-M)**. **(N-O)** Action potential threshold and rheobase of cells at RMP. M = male; f = female; ado = adolescent. Data are shown as mean +/− SEM. *p<0.05. **p<0.01. n=11 mice/sex/treatment for fMRI and n=6-7 mice/sex/treatment for electrophysiology.

### 3.4 Effects of adolescent binge drinking on functional connectivity and pyramidal cell excitability of the infralimbic cortex

Due to the infralimbic cortex’s known role in substance use related phenotypes [23,24] and our whole-brain connectivity data, we generated a seed-based map of the infralimbic cortex (**Fig 4A-B, Supp. Fig. 7**). In females, adolescent alcohol consumption increased RSFC between the infralimbic cortex and the primary motor area, agranular insular area, and olfactory tubercle (**Supp. Fig. 8**). To evaluate a possible cellular mechanism underlying connectivity changes, we tested passive properties and intrinsic excitability of pyramidal neurons in layer 2/3 of the infralimbic cortex (**Fig 4C-E**).

After adolescent alcohol consumption, the capacitance of infralimbic pyramidal cells was reduced (F_1,36_ = 5.8, p = 0.0217; **Fig 4F**), but there was no effect of treatment on membrane resistance (**Fig 4G**). A significant interaction of adolescent alcohol consumption and sex was detected for resting membrane potential (F_1,36_ = 8.9, p = 0.0050; **Fig 4H**), and Fisher’s LSD revealed that RMP was depolarized only in females (t_36_ = 3.304, p=0.0022) after adolescent alcohol consumption. The time constant did not vary across groups (**Fig 4I**). Action potential width was faster in males than in females (F_1,36_ = 5.9, p = 0.0198; **Fig 4J**), though this was not impacted by treatment. A three-way ANOVA of current-induced action potential firing revealed an (expected) effect of current (F_2,74_ = 181.4, p < 0.0001) and an interaction between adolescent alcohol consumption, current, and sex (F_2,74_ = 6.6, p = 0.0020; **Fig 4K**). Follow-up two-way ANOVA within sexes revealed that adolescent alcohol consumption had no effect in females (**Fig 4L**) but increased firing in males (F_1,18_ = 8.0, p = 0.0113; **Fig 4M**). An expected interaction of current and adolescent alcohol consumption was also seen in males (F_2,34_ = 5.4, p = 0.0100). Post hoc analyses within sexes revealed that as expected, current had a significant impact on firing in both sexes (female: F_2,38_ = 93.3, p < 0.0001; male: F_2,34_ = 89.1, p < 0.0001). Additional analyses of action potential threshold, rheobase, and current-induced firing at –70 mV did not reveal significant effects of adolescent alcohol (**Fig 4N-O, Supp. Fig. 9**). These results support both basal sex differences and an overall sex-specific effect of adolescent alcohol consumption on functional connectivity and intrinsic excitability of the infralimbic cortex, where only females show an increase in functional connectivity but both males and females show complex cellular adaptations.

### 3.5 Effects of adolescent binge drinking on functional connectivity and pyramidal cell excitability of the anterior insular cortex

Like with the infralimbic cortex, we conducted further follow-up analyses and experiments in the anterior insular cortex due to its documented role in substance use phenotypes [25]. A seed-based map of the anterior insular cortex did not reveal changes in functional connectivity due to alcohol (**Fig 5A-B**). In contrast to findings in the infralimbic PFC, no significant effects of adolescent alcohol consumption were found on RSFC of the anterior insular cortex with other ROIs (**Supp. Fig. 10**). Follow-up experiments examining *ex vivo* pyramidal cell passive properties and intrinsic excitability after adolescent alcohol exposure were conducted (**Fig 5C-E**). Adolescent alcohol consumption had no effect on the capacitance, membrane resistance, resting membrane potential, or time constant of anterior insula pyramidal neurons (**Fig 5F-I**). Action potential width was greater in adolescent alcohol-exposed mice (F_1,38_ = 6.9, p = 0.0125) and lower in males than in females (F_1,38_ = 4.2, p = 0.0465; **Fig 5J**). Current-induced firing at resting membrane potential was not different between groups (**Fig 5K-M**). Action potential threshold and rheobase at resting membrane potential were similar across groups (**Fig 5N-O**). Additional properties at –70mV were unaffected by alcohol (**Supp. Fig. 9**). In summary, only minor effects of adolescent alcohol consumption were found in the anterior insular cortex, providing contrast to pronounced alterations seen in the infralimbic cortex.

### 3.6 Effects of adolescent binge drinking on pyramidal cell excitability of the basolateral amygdala

We conducted identical experiments in the basolateral amygdala as a non-cortical comparison region known to be involved in substance use and stress [26,27]. Across most electrophysiological measurements, no significant differences due to alcohol were seen (**Supp. Fig. 11**), highlighting regional specificity of adolescent alcohol-induced brain perturbations.

## 4 DISCUSSION

Our current work sought to build upon a robust body of human and rodent literature suggesting adolescent alcohol consumption causes profound behavioral and neurobiological changes that last into adulthood, and that this effect differs across sexes. Using models of alcohol consumption and anxiety-like and exploratory behaviors, *in vivo* neuroimaging, and *ex vivo* cellular physiology, we demonstrate comprehensive changes in behavior and physiology following binge drinking. Overall, our data shows an interesting dichotomy between behavioral changes and physiological changes: while females showed the clearest adolescent alcohol-related behavioral and functional connectivity alterations, males had the clearest adolescent-alcohol induced infralimbic physiological adaptation (with more modest changes in behavior). This suggests that adolescent alcohol exposure produces qualitatively sexually divergent brain states in adulthood. Specifically, adolescent alcohol consumption promoted preference for alcohol and quinine-adulterated alcohol in adult females, and additional OFT and EPM tests revealed complementary sex-specific and alcohol dose-sensitive changes in exploration. fMRI experiments revealed alterations in whole brain resting state functional connectivity, with general downregulation of brain-wide connectivity in females, and general increased brain-wide connectivity in males. Patch clamp electrophysiology revealed region– and sex-specific changes in pyramidal neuron basal cellular properties and intrinsic excitability due to adolescent alcohol. Most notably, while physiological changes associated with increased infralimbic excitability were observed in both males and females, only females exhibited upregulated resting state functional connectivity in the infralimbic cortex. Cumulatively, these data provide evidence of long-term disruptions in addiction-related behaviors and brain function attributable to adolescent alcohol exposure, that manifest differently in the two sexes, and this dissociation between behavioral and neural phenotypes suggests that additional sex-dependent compensatory or circuit-level mechanisms remain to be identified.

A notable long-term consequence of adolescent alcohol exposure emerged in female adult alcohol preference, including when ethanol was adulterated with quinine. This has translational implications for human alcohol use, as a key component of human AUD is the continued use of alcohol despite negative consequences [28]. Notably, we and others have consistently observed that lower adolescent alcohol consumption corresponds to increased adult alcohol consumption [16]. This may suggest that adult alcohol consumption has an inverted U-shaped response to adolescent alcohol consumption. Importantly, it is well documented that female rodents often show higher baseline alcohol consumption [29,30]. Thus, adolescent exposure may amplify or alter an existing sexually-dimorphic adult alcohol preference. Additionally, we report that adolescent alcohol exposure increased exploration in males while decreasing exploration in females, and this scaled with volume of adolescent alcohol consumed. While adolescent alcohol exposure did not produce a generalized locomotor phenotype, adolescent alcohol dose was associated with lower adult open-arm exploration across several convergent EPM measures in females. The absence of corresponding OFT and EPM distance effects suggests that this phenotype is not readily explained by reduced locomotor capacity, which can be interpreted as an altered open-space exploration/avoidance or risk assessment. Overall, female-specific decreases in exploration coupled with female-specific susceptibility to aversion-resistant drinking accumulate to indicate that females with a history of adolescent alcohol use may be vulnerable to substance use through multiple mechanisms. Taken together, the quinine-choice and EPM results raise the possibility that adolescent alcohol alters how females evaluate reward under competing aversive or uncertain conditions, a hypothesis to be further explored in future work, and sheds light on the growing rate of AUD diagnoses in women [31]. Additional experiments further highlight that these effects are unique to the adolescent period, as adulthood alcohol exposure does not produce similar behavioral adaptations in the EPM.

Developmental imaging studies in humans suggest that adolescent alcohol modifies ongoing network maturation in a sex– and network-dependent manner [32–34]. To create synergy with, and greater possibilities for translation between, the human literature, whole-brain resting state fMRI was used to identify changes in neural circuits underlying changes in alcohol preference and exploration. We observed that females showed decreased, and males showed increased, brain-wide functional connectivity following adolescent binge drinking. We focused our attention and follow-up experiments on the infralimbic cortex, anterior insular cortex, and basolateral amygdala due to the robust literature supporting the relevance of these regions in alcohol phenotypes. For example, adolescent binge drinking is associated with differential engagement of the ventromedial PFC (similar to the mouse infralimbic cortex), insula, and amygdala during addiction-relevant decision-making tasks [35]. Separately, reduced engagement of the ventromedial PFC in a go/no-go task predicted increased substance use in a cohort of teenagers [36]. Our analysis supports the most pronounced functional deficits due to alcohol consumption in the infralimbic PFC. Specifically, females show increased functional connectivity, and both sexes showed altered measures of *ex vivo* pyramidal excitability, in the infralimbic cortex. In contrast, few differences are observed in the anterior insular cortex, and basolateral amygdala. This aligns with a larger body of literature demonstrating the unique sensitivity of the developing PFC to external insults such as alcohol.

Among the regions examined with cellular resolution, the infralimbic cortex showed the strongest evidence for persistent adaptation. Adolescent alcohol reduced membrane capacitance and produced sex-dependent effects on resting membrane potential and current-evoked firing. In males, increased action potential firing was observed when cells were held at their RMP, but not at −70 mV, suggesting a state-dependent alteration of the infralimbic cortex. Infralimbic cellular and functional connectivity deficits are known to be implicated in addiction-relevant behavioral aberrations. Gómez and colleagues [37] evaluated the impact of intragastric adolescent alcohol exposure on an attentional set shifting task and brain functional connectivity within the same rats. Within subjects, severity of functional connectivity deficits of multiple regions including the PFC were associated with reversal learning deficits, which were most pronounced in females. Thus, there is a clear link between adolescent alcohol exposure’s effects on functional connectivity and behavioral flexibility. Female-specific impairments in behavioral flexibility might also manifest as or be related to increased aversion-resistant drinking in adulthood. While aversion-resistant drinking is not a traditional measure of behavioral flexibility, prior literature has demonstrated a genetic link between behavioral flexibility and alcohol consumption [38].

Importantly, our preclinical rodent model data support other preclinical and human imaging findings, providing a framework of how the adolescent brain matures over development and responds to substances of abuse [6,13,39].

Males and females showed divergent effects when it came to *in vivo* versus *ex vivo* evaluations of the infralimbic cortex. While both sexes showed altered intrinsic-excitability phenotypes, these changes only paralleled the direction of female infralimbic functional connectivity alterations. While BOLD connectivity primarily reflects coordinated hemodynamic fluctuations, the corresponding *ex vivo* electrophysiological phenotype could be a candidate component of the circuit adaptation underlying changes in functional connectivity. One potential interpretation of this result in males is that *ex vivo* increased infralimbic excitability reflects a compensatory PFC adaptation, that is balanced via other mechanisms, resulting in neutral functional connectivity changes. An additional wholistic hypothesis is that this *ex vivo* neural adaptation in males is also maladaptive, but in behavioral domains not captured by the present assays, potentially including reversal learning or extinction tasks.

To summarize, voluntary binge-like alcohol consumption during adolescence produced persistent and sexually divergent adult adaptations across alcohol-related behavior, exploratory behavior, whole-brain functional connectivity, and PFC cellular physiology. We found that adolescent binge-like alcohol consumption led to increased aversion-resistant drinking in females, sex-specific changes in exploratory behaviors, and changes in whole brain functional connectivity that correspond to *ex vivo* cellular deficits. These data suggest that females may be more vulnerable to lasting addiction-related behavioral consequences of adolescent alcohol exposure, and these may be driven by brain-wide changes in functional connectivity. This is consistent with human data demonstrating that the positive relationship between adolescent binge drinking and later life alcohol use is dramatically stronger in females than in males [3]. This study provides biological targets for future studies evaluating the growing risk of alcohol use and misuse in women.

## 5 FUNDING

This study was supported by the National Institutes of Health [F32AA031396 (L.S.), F31AA030455 (A.S.), P50AA017823 (N.C.), R01AA029403 (N.C), R01AA031472 (N.C. and N.Z.), R01NS07816 (P.J.D.), T32NS115667 (M.S.H.), and T32GM154124 (Y.X.)], the Republic of Türkiye Ministry of National Education Scholarship (H.S.Ü.), the American Heart Association [AHA 24PRE1201066 (M.S.H.)], and the Pennsylvania State University [Student Engagement Network Grant (K.T.), Multi-Campus Research Experience for Undergraduates Fellowship (K.T.), Summer Undergraduate Research Experience Fellowships (B.J.D., E.M.), and Penn State University Eberly College of Science and The Huck Institutes of the Life Sciences Endowment Funds (N.C)].

## 6 DECLARATION OF COMPETING INTEREST

The authors declare that they have no competing interests.

## 7 ACKNOWLEDGEMENTS

We acknowledge the Huck Institutes of the Life Sciences High Field Magnetic Resonance Imaging Core Facility (RRID:SCR_024461) for use of the Bruker Biospec 70/30 and we thank the facility director Thomas Neuberger for help on the imaging protocol and in the facility. We thank Sophie Anderlind, Varya Zlotnik, and Isabella Girgis for their technical assistance.

## 8 CONTRIBUTIONS

LRS and NAC designed experiments. LRS, HSÜ, YX, ARS, KRG, KT, BJD, and EPM conducted experiments. LRS, HSÜ, YX, MSH, and LNB contributed to analysis. LRS and NAC wrote the manuscript with input from all authors. LRS, HSÜ, YX, MSH, ARS, KT, BJD, EM, NAC, NZ, and PJD acquired funding.

## Notes

### Competing Interest Statement

The authors have declared no competing interest.

### Summary of Updates

Updated experiments (slice electrophysiology) and new analyses of existing fMRI data

