## Supplemental Materials and Methods for "Adolescent alcohol consumption differentially alters sexually-divergent behaviors and infralimbic cortex function in mice"

***Running title: adolescent alcohol behavior and imaging***

**Seemiller et al.**

**Supplementary Materials**

**Supplementary Methods:**

*Sample sizes*

Experimental subjects were assigned to groups at random. Experimenters were blind to adolescent treatment group during adult testing. Outliers for adult phenotypes were identified in the ROUT outlier test (Q=1%), and mice had to be outliers for the majority of measures taken within the same test to be excluded. No mice met these criteria. Sample sizes were chosen based on preliminary studies and prior literature:

- Cohort 1 (2BC): 10 female H2O, 10 female EtOH, 9 male H2O, 11 male EtOH
- Cohort 2 (OFT, EPM): 10 female H2O, 10 female EtOH, 9 male H2O, 10 male EtOH
- Cohort 3 (fMRI): 11 female H2O, 11 female EtOH, 11 male H2O, 11 male EtOH
- Cohort 4 (EPM, ephys):
  - Infralimbic: 8 mice female H2O, 7 female EtOH, 8 male H2O, 9 male EtOH
  - BLA: 7 female H2O, 7 female EtOH, 7 male H2O, 7 male EtOH
  - AIC: 6 female H2O, 7 female EtOH, 6 male H2O, 6 male EtOH

*Drinking data analysis*

Drinking values that exceeded 2 g within 2 hr (roughly 100 g/kg for a 20 g mouse) were categorized as leaks and removed from drinking analyses. This drinking outlier threshold is consistent with our prior work (Sicher et al. 2024) that found consistent convergence in outliers identified through this method and the ROUT outlier method. These occasional missing data points (< 2% of DID data) due to bottle leakage were omitted in most drinking analyses and imputed as the average of related drinking values in analyses of total amounts consumed.

*Extended behavioral methods*

For two-bottle choice (2BC) testing, on PND 77-80, mice began acclimation to sipper tubes containing only H_2_O (tap water). Sipper tubes were weighed and refilled every 48-72 hr to monitor intake. Approximately 30+ days after DID (PND 84-87) mice underwent a continuous access 2BC where they received access to one tube of H_2_O and one tube of increasing concentrations of EtOH in H_2_O (3% w/v EtOH for 2 days, 7% w/v EtOH for 5 days, and 10% w/v EtOH for two weeks) which we have shown to reliably produce EtOH preference^16,17^. After this acclimation procedure, the 10% w/v EtOH in tap water was mixed with increasing concentrations of quinine hydrochloride (0.03, 0.1, 0.3, and 1 mM). Positions of H_2_O and EtOH tubes were alternated after every refill to avoid side preferences. Preference for EtOH solutions was defined as (EtOH consumed / total fluid consumed). Mice were weighed once weekly.

OFT and EPM occurred during the dark phase of the light cycle and under red light (8:00 AM - 4:00 PM). Mice were transferred to a procedure room and acclimated for one hr prior to behavior testing. First, mice were placed in the corner of an open field box (50 × 50 cm; with 20 cm black plexiglass walls) and allowed to explore for 5 min^18–20^. After OFT testing, mice were returned to home cages in the procedure room and allowed to acclimate again for at least one hr before EPM testing. Then, mice were placed in the center of an elevated plus maze (30 × 5 cm arms; 20 cm clear plexiglass walls on closed arms; maze elevated 40 cm) and allowed to explore for 5 min^18^. For mice used in electrophysiology experiments, only EPM (and not OFT) was performed 30 days after DID. EPM data were excluded after mice occasionally fell off of the maze (three times for primary behavior experiment).

OFT and EPM experiments were video-recorded with a camera at a frame rate of 33.6 frames per second. Movement was tracked using DeepLabCut (V2.2.1). The head, middle of body, and trunk of the tail were tracked throughout experiments. Tracking employed transfer learning to retrain residual network EfficientNet-B0 (effnet_b0). Training was sufficient when confidence in the position was >99%. Data were exported as a CSV file and analyzed in MATLAB.

Across both OFT and EPM tests, the Euclidean distance of the middle of the body between frames was classified as the distance traveled. The number of frames that a mouse was in the polygon bounded by the zone of interest was used to calculate time spent in a zone of interest. Pixels/frame was converted to cm/s using known arena measurements and camera frame rate. Randomly selected videos were hand-scored and used to validate automated behavior analysis.

*OFT and EPM 24 hr after adolescent DID (Methods for supplemental Figure 3)*

Male and female C57BL/6J mice were bred in-house, weaned and moved into single housing on a reverse light cycle on PND 21, and underwent DID PND 29-53 (+/- 1 day). One day after the last alcohol access session and the conclusion of DID (PND 53 +/- 1), mice underwent OFT and EPM testing. One subject jumped out of the OFT box during testing, so OFT time in center and distance traveled were not collected for that mouse. Statistical analysis and outlier testing was done as described in the main text. No outliers were detected for OFT or EPM data sets.

*OFT and EPM 30 days after adult DID (Methods for supplemental Figure 4)*

For supplemental adult experiments, male and female C57BL/6J mice were ordered from Jackson Laboratory (Bar Harbor, ME, USA). At least one week prior to DID, subjects were moved and single housed in a reversed light cycle room. DID was conducted as described in the main text, except that it occurred during adulthood (PND 62-86). OFT and EPM testing began on PND 118 to mimic the 30 day washout in adolescent experiments. Statistical analysis and outlier testing was performed as described in the main text. No mice fell off the EPM, but one outlier subject was removed from EPM analysis.

*Extended fMRI methods*

*fMRI acclimation and data acquisition*: Mice were acclimated to the imaging environment for 4 days prior to testing. Briefly, mice were anesthetized with 3% isoflurane while their limbs were lightly bound using surgical tape. Mice were gently positioned in a cylindrical restrainer tube and secured with a bite bar. After mice awoke from anesthesia (assessed by recovery of awake respiration rate, usually 10-20 mins), they were monitored while acclimating for increasing amounts of time. Specifically, mice were restrained for 15, 30, 45, and 60 min on acclimation days 1-4, respectively, while a soundtrack of fMRI noises was played. Afterward, on PND 83-88 (at least 30 days after DID), awake resting state imaging was then performed using a 7T MRI system and Bruker Console (Billerica, MA) using Paravision 7.0.0. Again, mice were briefly anesthetized with 3% isoflurane while being restrained. Once restrained, anesthesia was stopped and mice were allowed to wake up before imaging. A gradient-echo echo-planar imaging (GE-EPI) sequence for T2*-weighted resting-state scans was used for data collection (repetition time (TR) = 1.5 s; echo time (TE) = 15 ms; field of view (FOV) = 1.6 × 1.6 cm; image matrix size 64 x 64; slice number = 16; slice thickness 0.75 mm; slice gap = 0 mm). Each scan lasted 10 min and collected 400 volumes, with three scans per mouse. Rapid imaging with refocused echoes (RARE) was used to acquire T1-weighted anatomical images (TR = 1.5 s; TE = 8 ms; FOV = 1.6 × 1.6 cm; image matrix size = 128 × 128; slice number = 16; slice thickness = 0.75 mm; RARE factor = 8; repetition number = 6; slice gap = 0 mm).

*fMRI image preprocessing:* Framewise displacement (FD) was calculated to quantify motion (Power et al. 2012): FD = ∣ Δx ∣ + ∣ Δy ∣ + ∣ Δz ∣ + r * (∣ Δα ∣ + ∣ Δβ ∣ + ∣ Δγ ∣). r = 2 mm represents the length from the cortex to the center of the brain. Δx, Δy, and Δz (translation distances) and Δα, Δβ, and Δγ (rotations) across x, y, and z axis were calculated using "imregtform" geometric transformation function in MATLAB version R2023b (The MathWorks Inc., Natick, MA, United States). Volumes exhibiting motion over half of the in-plane voxel size (0.125mm) and their temporally adjacent volumes were removed from analysis. Data were manually co-registered to a reference template for anatomical assessment and normalized to Allen Mouse Brain Atlas (Lein et al. 2007) using a custom-developed MATLAB program (Liu et al. 2020). Motion calculation and correction were performed using the Statistical Parametric Mapping (SPM12) package (http://www.fil.ion.ucl.ac.uk/spm/). We performed spatial smoothing using a Gaussian kernel (full-width-half-maximum = 0.375mm, 1.5x the x-y plane voxel size). We performed nuisance regression to eliminate signals originating from white matter (WM) and ventricle voxels, thereby reducing physiological noise (respiration, heart rate, etc.), along with regressing out the six motion parameters for each fMRI frame. Finally, a band-pass filter (0.01-0.1 Hz) was used. For consistency across data sets, 350 volumes from each scan (originally containing 400 volumes) were used.

*fMRI image postprocessing:* A total of 62 regions of interest (ROIs) were defined based on definitions from the Allen Mouse Brain Atlas (Lein et al. 2007). ROIs were assigned to 10 brain systems: hippocampal region, retrohippocampal region, striatum, sensory-motor cortex, polymodal association cortex, olfactory cortex, pallidum, thalamus, hypothalamus, and midbrain. Resting state functional connectivity between brain regions was analyzed using an ROI-based approach specifically testing Pearson correlations between the average activity time course between each ROI pair. Functional connectivity matrices were calculated using Fisher’s z-transformed Pearson’s correlation coefficients within each mouse. Seed-based functional connectivity analysis was also used to directly determine regions with strong functional connectivity with selected regions. Three scans were collected per mouse, with each scan lasting 10 minutes. After applying motion scrubbing and excluding scans with more than 10% of volumes discarded, all animals retained at least one scan that could be included in the analysis. The total number of usable scans per group was as follows: alcohol female (11 mice): 32 scans, alcohol male (11 mice): 33 scans, control female (11 mice): 28 scans, and control male (11 mice): 30 scans.

*Extended ex vivo electrophysiology methods*

Mice were anesthetized with inhaled isoflurane and rapidly decapitated. Brains were rapidly extracted and immediately placed in ice-cold N-methyl-D-glucamine (NMDG) solution containing the following (in mM): 93 NMDG, 2.5 KCl, 1.2 NaH2PO4, 30 NaHCO3, 20 HEPES, 25 dextrose, 5 ascorbic acid, 2 thiourea, 3 sodium pyruvate, 10 MgSO4-7H2O, 0.5 CaCl2-2H2O, 306-310mOsm, pH 7.4 (Ting et al., 2018). 300 uM Coronal sections containing the infralimbic cortex, anterior insula, or basolateral amygdala were prepared with a Compresstome vibrating microtome (Precisionary Instruments). Slices rested in oxygenated NMDG cutting solution (31°C) for a maximum of 8 min then were transferred to oxygenated artificial cerebrospinal fluid (aCSF) at 31°C, containing the following (in mM): 124 NaCl, 4.0 KCl, 1.2 MgSO4-7H2O, 2.0CaCl2-2H2O, 1 NaH2PO4, 25.95 NaHCO3, 9.99 dextrose, 305-308mOsmo. Slices rested in oxygenated aCSF (31°C) for at least 1 hr prior to transfer to a submerged chamber for experiments. In the submerged chamber, slices were perfused with heated, oxygenated aCSF at a rate of 2 mL/min. Recording electrodes (3-5 MΩ) were pulled from thin-walled borosilicate glass capillaries using a Narishige PC-100 vertical puller. Electrodes for all experiments were filled with a potassium-gluconate based recording solution (in mM): 135 potassium gluconic acid, 5 NaCl, 2 MgCl2-6H2O, 10 HEPES, 0.6 EGTA, 4 Na2ATP, 0.4 Na2-GTP, 287-290 mOsm, pH 7.35.

Following 5 min of baseline recording to allow the internal to diffuse in the cell, a series of depolarizing ramp currents (100 pA over 1 second) were injected to determine rheobase. Current-induced action potential firing was assessed using discrete steps of hyperpolarizing to depolarizing current (-100 to 350 pA in 10 pA steps, each step lasting 250 ms). Experiments were performed at each neuron’s resting membrane potential (RMP) then repeated at a common holding potential of -70 mV. Cells were discarded from analysis if they had a resting membrane potential > -45 mV, tonically fired action potentials, did not fire action potentials in response to current steps, or had an access resistance exceeding 40 MΩ at the end of the experiment.

**Supplementary Results:**

Daily alcohol consumption was analyzed via two-way (sex, day) mixed model ANOVA and body weights were analyzed via three-way (treatment, sex, week) mixed model ANOVA. In the first experimental cohort, Cohort 1, adolescent females consumed more alcohol than adolescent males (F_1,19_ = 10.2, *p* = 0.0048). An interaction between treatment and sex was detected for body weight (F_1,33_ = 4.1, *p* = 0.0498), and post-hoc two-way ANOVA (treatment, week) within sexes revealed that alcohol-treated females weighed less than female water controls (F_1,18_ = 4.5, *p* = 0.0476) and alcohol exposure interacted with week (F_3,54_ = 3.2, *p* = 0.0309). No treatment effect was seen for male body weights. Treatment effects were not detected for remaining Cohorts 2-4.

During adolescent DID, alcohol consumption across all cohorts was affected by day (Cohort 1: F_6,106_ = 8.1, p < 0.0001; Cohort 2: F_7,129_ = 11.3, p < 0.0001; Cohort 3: F_15,150_ = 10.8, p < 0.0001; Cohort 4: F_6,101_ = 11.5, p < 0.0001). Females weighed less than males (Cohort 1: F_1,33_ = 37.4, p < 0.0001; Cohort 2: F_1,36_ = 42.2, p < 0.0001; Cohort 3: F_1,40_ = 71.6, p < 0.0001; Cohort 4: F_1,38_ = 48.7, p < 0.0001) and body weights changed across weeks (Cohort 1: F_2,58_ = 214.5, p < 0.0001; Cohort 2: F_2,76_ = 289.8, p < 0.0001; Cohort 3: F_2,77_ = 462.8, p < 0.0001; Cohort 4: F_2,72_ = 132.0, p < 0.0001). Body weights were also significantly impacted by week in the sex-specific analysis of cohort 1 (female: F_2,43_ = 300.1, p < 0.0001; male: F_2,24_ = 62.8, p < 0.0001). Further, body weights were influenced by an interaction of week and sex in cohorts 1-3 (Cohort 1: F_3,99_ = 3.1, p = 0.0308; Cohort 2: F_3,108_ = 3.8, p = 0.0129; Cohort 3: F_3,120_ = 8.9, p < 0.0001). *Data are represented in supplemental* ***Fig 1***.

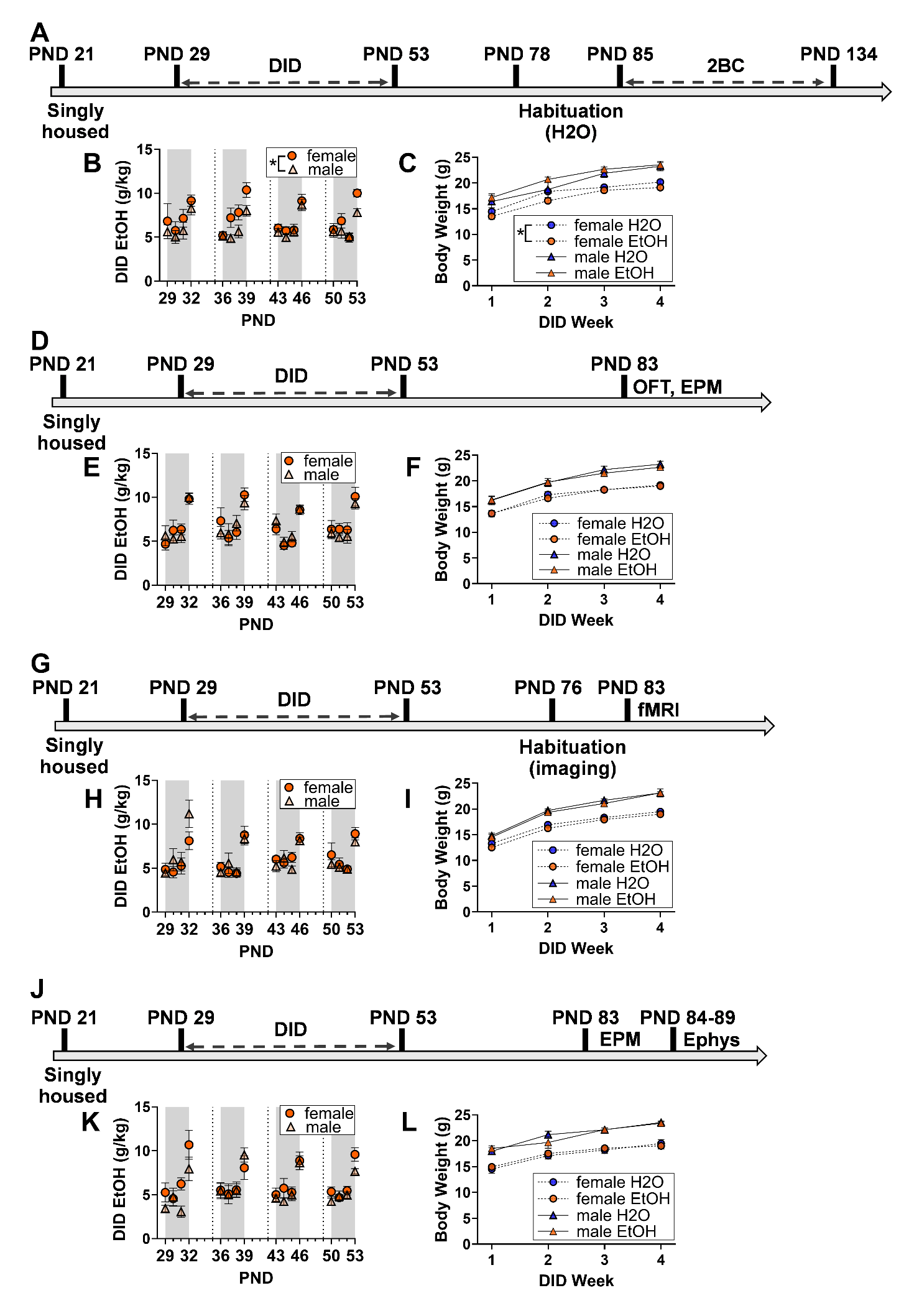
**Supp Figure 1:** Experimental timelines, adolescent alcohol consumption, and body weights across cohorts. Four separate experiments tested the influence of adolescent drinking in the dark (DID) on adult aversion-resistant alcohol preference via two-bottle choice (2BC; **A-C**), exploratory behavior in the open field test (OFT) and elevated plus maze (EPM; (**D-F**), whole brain resting state functional connectivity (**G-I**), and pyramidal cell intrinsic excitability (**J-L**). PND is approximate (see methods). Data are shown as mean +/- SEM. **p* < 0.05. n = 9-11 (**A-C**), 8-10 (**D-F**), 11 (**G-I**), and 6-9 (**J-L**) per sex per treatment.

| **Dependent variable** | **Effect** | **F (DFn, DFd)** | **p** |
| --- | --- | --- | --- |
| Ado DID EtOH – experiment 1 (Fig 1B)  2-way (Day, Sex) ANOVA | Day | F (5.648, 105.8) = 8.129 | <0.0001 |
|  | Sex | F (1, 19) = 10.21 | 0.0048 |
|  | Day x Sex | F (15, 281) = 0.6743 | 0.8092 |
| Body weight – experiment 1 (Fig 1C)  3-way (Week, Sex, Treatment) ANOVA | Week | F (1.757, 57.99) = 214.5 | <0.0001 |
|  | Sex | F (1, 33) = 37.41 | <0.0001 |
|  | Treatment | F (1, 33) = 0.02298 | 0.8804 |
|  | Week x Sex | F (3, 99) = 3.083 | 0.0308 |
|  | Week x Treatment | F (3, 99) = 0.3734 | 0.7724 |
|  | Sex x Treatment | F (1, 33) = 4.145 | 0.0498 |
|  | Week x Sex x Treatment | F (3, 99) = 2.443 | 0.0686 |
| Body weight – experiment 1 (Fig 1C)  Post-hoc within males  2-way (Week, Treatment) ANOVA | Week | F (1.597, 23.95) = 62.79 | <0.0001 |
|  | Treatment | F (1, 15) = 1.070 | 0.3172 |
|  | Week x Treatment | F (3, 45) = 0.9191 | 0.4393 |
| Body weight – experiment 1 (Fig 1C)  Post-hoc within females  2-way (Week, Treatment) ANOVA | Week | F (2.386, 42.96) = 300.1 | <0.0001 |
|  | Treatment | F (1, 18) = 4.520 | 0.0476 |
|  | Week x Treatment | F (3, 54) = 3.188 | 0.0309 |
| Ado DID EtOH – experiment 2 (Fig 1E)  2-way (Day, Sex) ANOVA | Day | F (7.433, 128.8) = 11.26 | <0.0001 |
|  | Sex | F (1, 18) = 0.1146 | 0.7389 |
|  | Day x Sex | F (15, 260) = 0.5982 | 0.8754 |
| Body weight – experiment 2 (Fig 1F)  3-way (Week, Sex, Treatment) ANOVA | Week | F (2.100, 75.58) = 289.8 | <0.0001 |
|  | Sex | F (1, 36) = 42.17 | <0.0001 |
|  | Treatment | F (1, 36) = 0.2492 | 0.6207 |
|  | Week x Sex | F (3, 108) = 3.765 | 0.0129 |
|  | Week x Treatment | F (3, 108) = 0.4191 | 0.7396 |
|  | Sex x Treatment | F (1, 36) = 0.002849 | 0.9577 |
|  | Week x Sex x Treatment | F (3, 108) = 1.094 | 0.3549 |
| Ado DID EtOH – experiment 3 (Fig 1H)  2-way (Day, Sex) ANOVA | Day | F (15, 150) = 10.78 | <0.0001 |
|  | Sex | F (1, 10) = 0.0008475 | 0.9773 |
|  | Day x Sex | F (15, 145) = 1.231 | 0.2553 |
| Body weight – experiment 3 (Fig 1I)  3-way (Week, Sex, Treatment) ANOVA | Week | F (1.916, 76.63) = 462.8 | <0.0001 |
|  | Sex | F (1, 40) = 71.57 | <0.0001 |
|  | Treatment | F (1, 40) = 1.597 | 0.2136 |
|  | Week x Sex | F (3, 120) = 8.902 | <0.0001 |
|  | Week x Treatment | F (3, 120) = 0.2274 | 0.8772 |
|  | Sex x Treatment | F (1, 40) = 0.1586 | 0.6926 |
|  | Week x Sex x Treatment | F (3, 120) = 0.3506 | 0.7888 |
| Ado DID EtOH – experiment 4 (Fig 1K)  2-way (Day, Sex) ANOVA | Day | F (5.844, 100.5) = 11.53 | <0.0001 |
|  | Sex | F (1, 18) = 1.707 | 0.2078 |
|  | Day x Sex | F (5.844, 100.5) = 1.292 | 0.2686 |
| Body weight – experiment 4 (Fig 1L)  3-way (Week, Sex, Treatment) ANOVA | Week | F (1.884, 71.61) = 132.0 | <0.0001 |
|  | Sex | F (1, 38) = 48.70 | <0.0001 |
|  | Treatment | F (1, 38) = 0.008463 | 0.9272 |
|  | Week x Sex | F (1.884, 71.61) = 1.649 | 0.2007 |
|  | Week x Treatment | F (1.884, 71.61) = 1.576 | 0.2149 |
|  | Sex x Treatment | F (1, 38) = 0.1947 | 0.6615 |
|  | Week x Sex x Treatment | F (1.884, 71.61) = 1.877 | 0.1627 |

**Supp Table 1**: All ANOVA outcomes for adolescent DID and body weight data presented in Supplemental Figure 1.

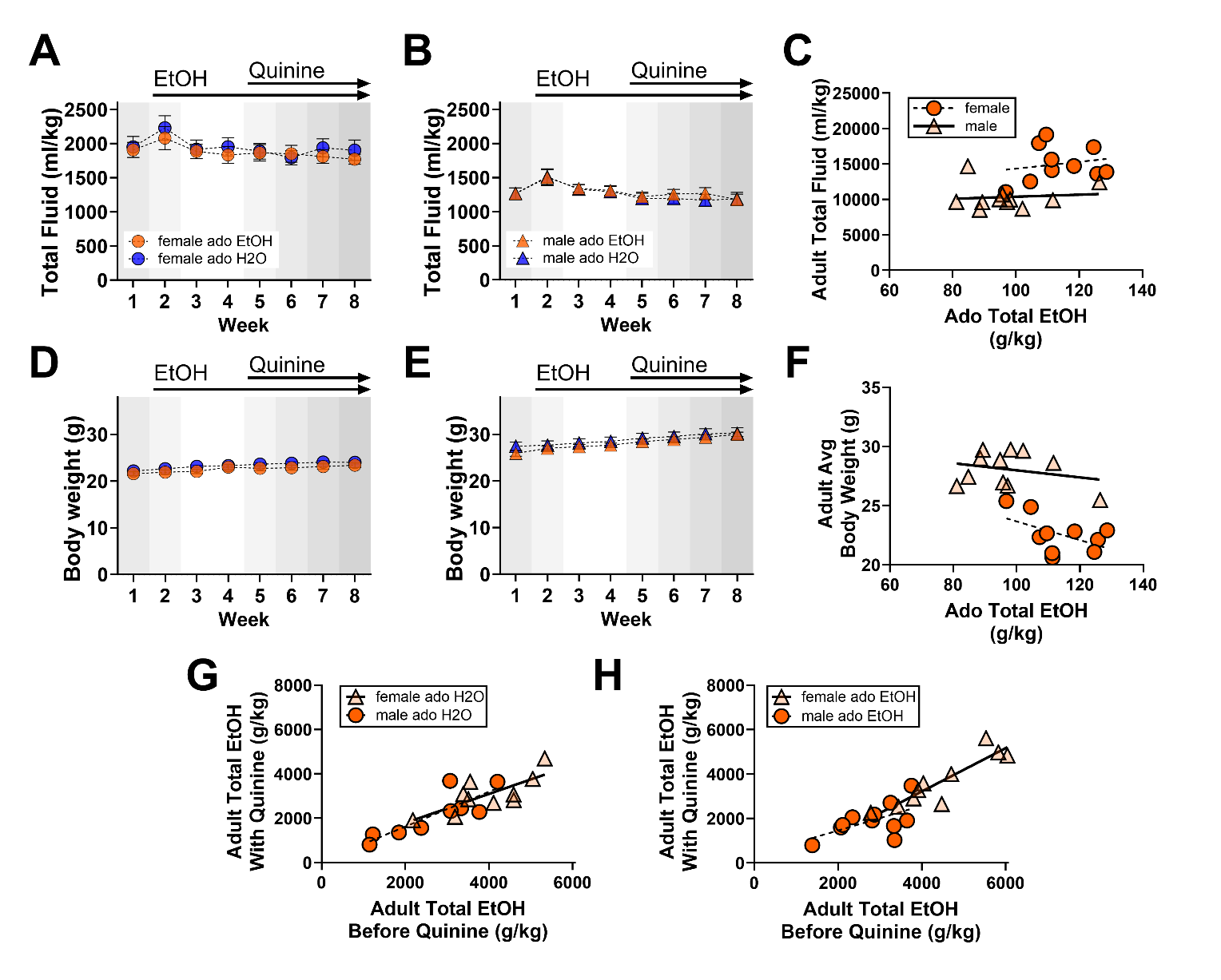

**Supp Fig 2.** Adolescent alcohol consumption did not affect total fluid consumption (**A-C**) or body weight (**D-F**) during two-bottle choice assessment of alcohol preference. Total alcohol consumed in the first weeks of adult two-bottle choice testing without quinine was significantly associated with total alcohol consumed with quinine (total EtOH no quinine: 0.6538 ± 0.1799, p = 0.0009), but this did not differ across sex or adolescent alcohol treatment groups (**G-H**). Ado = Adolescent. Week 1 represents consumption of H2O only, Week 2 includes 3-7% EtOH, Week 3-4 represents 10% EtOH, and Weeks 5-8 represent 10% EtOH with increasing concentrations of quinine (0.03, 0.1, 0.3, and 1mM). Plots labeled as “Before Quinine” represent data from Weeks 1-4, and plots labeled as “With Quinine” represent data from Weeks 5-8. In scatter plots, a simple linear regression was used to generate lines of best fit and aid interpretation but were independent from statistical analysis. Data are shown as mean +/- SEM. * *p* <0.05. n=9-11/sex/treatment.

| **Dependent variable** | **Effect** | **F (DFn, DFd)** | **p** |
| --- | --- | --- | --- |
| Female EtOH Preference (Fig 2A)  2-way (Week, Treatment) mixed effects ANOVA | Week | F (3.879, 69.82) = 109.1 | <0.0001 |
|  | Treatment | F (1, 18) = 4.688 | 0.0441 |
|  | Week x Treatment | F (6, 108) = 1.541 | 0.1717 |
| Female EtOH Consumption (Fig 2B)  2-way (Week, Treatment) mixed effects ANOVA | Week | F (4.501, 81.01) = 99.42 | <0.0001 |
|  | Treatment | F (1, 18) = 1.530 | 0.2320 |
|  | Week x Treatment | F (6, 108) = 1.562 | 0.1654 |
| Female H2O Consumption (Fig 2C)  2-way (Week, Treatment) mixed effects ANOVA | Week | F (4.165, 74.97) = 124.5 | <0.0001 |
|  | Treatment | F (1, 18) = 3.209 | 0.0900 |
|  | Week x Treatment | F (7, 126) = 1.991 | 0.0613 |
| Female Fluid Consumption (Supp Fig 1A)  2-way (Week, Treatment) mixed effects ANOVA | Week | F (3.979, 71.62) = 4.473 | 0.0028 |
|  | Treatment | F (1, 18) = 0.2025 | 0.6581 |
|  | Week x Treatment | F (7, 126) = 0.5347 | 0.8068 |
| Female Adult Body Weight (Supp Fig 1D)  2-way (Week, Treatment) mixed effects ANOVA | Week | F (4.113, 74.03) = 40.14 | <0.0001 |
|  | Treatment | F (1, 18) = 1.690 | 0.2099 |
|  | Week x Treatment | F (7, 126) = 1.471 | 0.1835 |
| Male EtOH Preference (Fig 2A)  2-way (Week, Treatment) mixed effects ANOVA | Week | F (3.197, 57.54) = 27.01 | <0.0001 |
|  | Treatment | F (1, 18) = 0.1918 | 0.6667 |
|  | Week x Treatment | F (6, 108) = 0.9424 | 0.4680 |
| Male EtOH Consumption (Fig 2B)  2-way (Week, Treatment) mixed effects ANOVA | Week | F (3.240, 58.32) = 43.65 | <0.0001 |
|  | Treatment | F (1, 18) = 0.01788 | 0.8951 |
|  | Week x Treatment | F (6, 108) = 0.8304 | 0.5489 |
| Male H2O Consumption (Fig 2C)  2-way (Week, Treatment) mixed effects ANOVA | Week | F (3.796, 68.32) = 39.85 | <0.0001 |
|  | Treatment | F (1, 18) = 0.1121 | 0.7417 |
|  | Week x Treatment | F (7, 126) = 0.7018 | 0.6705 |
| Male Fluid Consumption (Supp Fig 1A)  2-way (Week, Treatment) mixed effects ANOVA | Week | F (3.346, 60.23) = 8.415 | <0.0001 |
|  | Treatment | F (1, 18) = 0.05844 | 0.8117 |
|  | Week x Treatment | F (7, 126) = 0.3125 | 0.9472 |
| Male Adult Body Weight (Supp Fig 1D)  2-way (Week, Treatment) mixed effects ANOVA | Week | F (1.919, 34.55) = 41.94 | <0.0001 |
|  | Treatment | F (1, 18) = 0.5886 | 0.4529 |
|  | Week x Treatment | F (7, 126) = 1.018 | 0.4216 |

**Supp Table 2**: All ANOVA outcomes for experiment 1 examining adolescent DID effects on adult alcohol preference (presented in main text Figure 1 and Supplemental Figure 2).

| **Dependent variable** | **Distribution, Link** | **Effect** | **Estimate** | **SE** | **p** | **Coefficient Lower** | **Coefficient Upper** |
| --- | --- | --- | --- | --- | --- | --- | --- |
| Adult Average EtOH Preference Before Quinine (Fig 2G) | Normal, Identity | Sex (Male) | -1.22 | 0.51557 | 0.03011 | -2.3077 | -0.1322 |
|  |  | Ado EtOH | -0.0092989 | 0.0040242 | 0.033658 | -0.017789 | -0.00080853 |
|  |  | Sex x Ado EtOH | 0.0098572 | 0.0047175 | 0.052005 | -9.5843e-05 | 0.01981 |
| Adult Total EtOH (g/kg) Before Quinine (Fig 2H) | Normal, Identity | Sex (Male) | -7300.1 | 2334.3 | 0.006135 | -12225 | -2375.1 |
|  |  | Ado EtOH | -77.03 | 19.392 | 0.00098475 | -117.94 | -36.116 |
|  |  | Sex x Ado EtOH | 46.756 | 21.202 | 0.04149 | 2.0245 | 91.488 |
| Post-hoc Female Adult Total EtOH (g/kg) Before Quinine (Fig 2I) | Normal, Identity | Ado EtOH | 5.493 | 2.7744 | 0.063213 | -0.33576 | 11.322 |
| Post-hoc Male Adult Total EtOH (g/kg) Before Quinine (Fig 2I) | Normal, Identity | Ado EtOH | 1.9742 | 3.0892 | 0.53084 | -4.5161 | 8.4644 |
| Adult Total H2O (ml/kg) Before Quinine (Fig 2I) | Normal, Identity | Sex (Male) | 4185.9 | 3304.3 | 0.22231 | -2785.6 | 11157 |
|  |  | Ado EtOH | 41.761 | 24.241 | 0.10308 | -9.3835 | 92.906 |
|  |  | Sex x Ado EtOH | -42.641 | 30.437 | 0.17922 | -106.86 | 21.576 |
| Adult Average EtOH Preference After Quinine (Fig 2J) | Normal, Identity | Sex (Male) | -1.1404 | 0.55206 | 0.054451 | -2.3051 | 0.024371 |
|  |  | Ado EtOH | -0.010501 | 0.0043085 | 0.02608 | -0.019591 | -0.0014108 |
|  |  | Sex x Ado EtOH | 0.0090691 | 0.0050514 | 0.090393 | -0.0015884 | 0.019727 |
| Adult Total EtOH (g/kg) After Quinine (Fig 2K) | Normal, Identity | Sex (Male) | -7122.3 | 2759.8 | 0.019438 | -12945 | -1299.7 |
|  |  | Ado EtOH | -85.924 | 22.866 | 0.0015686 | -134.17 | -37.68 |
|  |  | Sex x Ado EtOH | 45.618 | 25.074 | 0.086519 | -7.2836 | 98.52 |
| Adult Total H2O (ml/kg) After Quinine (Fig 2L) | Normal, Identity | Sex (Male) | 6981.3 | 3163.4 | 0.041357 | 307.19 | 13655 |
|  |  | Ado EtOH | 73.618 | 23.207 | 0.005571 | 24.655 | 122.58 |
|  |  | Sex x Ado EtOH | -65.655 | 29.139 | 0.037746 | -127.13 | -4.1768 |
| Post-hoc Female Adult Total H2O (ml/kg) After Quinine (Fig 2L) | Normal, Identity | Ado EtOH | -6.1345 | 3.6053 | 0.10606 | -13.709 | 1.44 |
| Post-hoc Male Adult Total H2O (ml/kg) After Quinine (Fig 2L) | Normal, Identity | Ado EtOH | 4.9504 | 5.0169 | 0.33685 | -5.5898 | 15.491 |
| Adult Total Fluid (ml/kg) (Supp Fig 1C) | Normal, Identity | Sex (Male) | -1030 | 8088.2 | 0.90016 | -18095 | 16035 |
|  |  | Ado EtOH | -1.8252 | 64.541 | 0.97777 | -137.99 | 134.34 |
|  |  | Sex x Ado EtOH | -32.213 | 73.819 | 0.66806 | -187.96 | 123.53 |
| Adult Average Body Weight (g) (Supp Fig 1F) | Normal, Identity | Sex (Male) | -1.5712 | 5.7126 | 0.7866 | -13.624 | 10.481 |
|  |  | Ado EtOH | -0.080137 | 0.043694 | 0.084215 | -0.17232 | 0.01205 |
|  |  | Sex x Ado EtOH | 0.058422 | 0.052389 | 0.2803 | -0.05211 | 0.16895 |
| Adult Total EtOH (g/kg) After Quinine (Supp Fig 1G-H) | Normal, Identity | Sex (Male) | -430.45 | 874.5 | 0.62593 | -2211.8 | 1350.9 |
|  |  | Adult EtOH Before Quinine | 0.65376 | 0.17992 | 0.00096826 | 0.28727 | 1.0202 |
|  |  | Ado EtOH group | -1153.3 | 1037.1 | 0.27444 | -3265.8 | 959.32 |
|  |  | Sex x Adult EtOH Before Quinine | 0.1325 | 0.24655 | 0.5947 | -0.36971 | 0.63471 |
|  |  | Sex x Ado EtOH Group | 1451 | 1308.4 | 0.27571 | -1214.2 | 4116.2 |
|  |  | Adult EtOH Before Quinine x Ado EtOH Group | 0.31943 | 0.24175 | 0.19576 | -0.17299 | 0.81185 |
|  |  | Sex x Adult EtOH Before Quinine x Ado EtOH Group | -0.54941 | 0.36712 | 0.14431 | -1.2972 | 0.19838 |

**Supp Table 3**: All GLME outcomes for experiment 1 examining adolescent DID effects on adult alcohol preference (presented in main text Figure 1 and Supplemental Figure 2).

| **Dependent variable** | **Effect** | **F (DFn, DFd)** | **p** |
| --- | --- | --- | --- |
| OFT Latency to Enter Center (Fig 2A)  2-way (Sex, Treatment) ANOVA | Sex | F (1, 35) = 2.239e-007 | 0.9996 |
|  | Treatment | F (1, 35) = 0.03709 | 0.8484 |
|  | Sex x Treatment | F (1, 35) = 0.5840 | 0.4499 |
| OFT Time in Center (Fig 2B)  2-way (Sex, Treatment) ANOVA | Sex | F (1, 35) = 3.126 | 0.0858 |
|  | Treatment | F (1, 35) = 0.09244 | 0.7629 |
|  | Sex x Treatment | F (1, 35) = 0.02525 | 0.8747 |
| OFT Distance Traveled (Fig 2C)  2-way (Sex, Treatment) ANOVA | Sex | F (1, 35) = 0.3328 | 0.5677 |
|  | Treatment | F (1, 35) = 3.918 | 0.0557 |
|  | Sex x Treatment | F (1, 35) = 0.3545 | 0.5554 |
| EPM Latency to Enter Open Arm (Fig 2G)  2-way (Sex, Treatment) ANOVA | Sex | F (1, 32) = 1.083 | 0.3059 |
|  | Treatment | F (1, 32) = 1.212 | 0.2792 |
|  | Sex x Treatment | F (1, 32) = 4.369 | 0.0446 |
| EPM Time in Open Arms (Fig 2H)  2-way (Sex, Treatment) ANOVA | Sex | F (1, 32) = 0.01212 | 0.9130 |
|  | Treatment | F (1, 32) = 0.01757 | 0.8954 |
|  | Sex x Treatment | F (1, 32) = 0.7769 | 0.3847 |
| EPM Open Arm Entries (Fig 2I)  2-way (Sex, Treatment) ANOVA | Sex | F (1, 32) = 0.1890 | 0.6667 |
|  | Treatment | F (1, 32) = 0.02276 | 0.8810 |
|  | Sex x Treatment | F (1, 32) = 2.572 | 0.1186 |
| EPM Head Dips (Fig 2M)  2-way (Sex, Treatment) ANOVA | Sex | F (1, 32) = 0.01434 | 0.9054 |
|  | Treatment | F (1, 32) = 0.008887 | 0.9255 |
|  | Sex x Treatment | F (1, 32) = 0.02216 | 0.8826 |
| EPM Distance Traveled (Fig 2N)  2-way (Sex, Treatment) ANOVA | Sex | F (1, 32) = 0.4477 | 0.5082 |
|  | Treatment | F (1, 32) = 0.1038 | 0.7495 |
|  | Sex x Treatment | F (1, 32) = 0.06153 | 0.8057 |

**Supp Table 4**: All ANOVA outcomes for experiment 2 examining adolescent DID effects on adult exploratory behavior (presented in main text Figure 2). Post-hoc t tests following the interaction of sex and treatment detected for latency to enter an open arm of the EPM did not reveal significant differences between alcohol- and water-treatment groups within males (t=1.493, df=7.103, p=0.1784) or females (t=1.548, df=9.423, p=0.1546).

| **Dependent variable** | **Distribution, Link** | **Effect** | **Estimate** | **SE** | **p** | **Coefficient Lower** | **Coefficient Upper** |
| --- | --- | --- | --- | --- | --- | --- | --- |
| OFT Latency to Enter Center (Fig 2D) | Poisson, Log | Sex (Male) | -0.78722 | 0.94129 | 0.41529 | -2.7827 | 1.2082 |
|  |  | Ado EtOH | -0.011345 | 0.0061267 | 0.082619 | -0.024333 | 0.0016435 |
|  |  | Sex x Ado EtOH | 0.0091045 | 0.0086616 | 0.082619 | -0.0092572 | 0.027466 |
| OFT Time in Center (Fig 2E) | Normal, Identity | Sex (Male) | -11.867 | 13.605 | 0.39597 | -40.708 | 16.974 |
|  |  | Ado EtOH | -0.12627 | 0.084656 | 0.15526 | -0.30573 | 0.053192 |
|  |  | Sex x Ado EtOH | 0.050401 | 0.12437 | 0.69066 | -0.21325 | 0.31406 |
| OFT Distance Traveled (Fig 2F) | Normal, Identity | Sex (Male) | -268.7 | 1160.8 | 0.81987 | -2729.5 | 2192.1 |
|  |  | Ado EtOH | 4.8045 | 7.1446 | 0.51088 | -10.341 | 19.95 |
|  |  | Sex x Ado EtOH | 3.5844 | 10.673 | 0.74135 | -19.041 | 26.21 |
| EPM Latency to Enter Open Arm (Fig 2J) | Poisson, Log | Sex (Male) | -3.6773 | 2.8487 | 0.21628 | -9.7491 | 2.3945 |
|  |  | Ado EtOH | 0.046292 | 0.011457 | 0.0010677 | 0.021872 | 0.070712 |
|  |  | Sex x Ado EtOH | 0.021018 | 0.025064 | 0.41486 | -0.032404 | 0.07444 |
| EPM Time in Open Arms (Fig 2K) | Normal, Identity | Sex (Male) | -100.69 | 45.956 | 0.045874 | -199.25 | -2.1203 |
|  |  | Ado EtOH | -0.5524 | 0.28632 | 0.074214 | -1.1665 | 0.061697 |
|  |  | Sex x Ado EtOH | 0.98271 | 0.42438 | 0.036258 | 0.072505 | 1.8929 |
| Post-hoc Female EPM Time in Open Arms (Fig 2K) | Normal, Identity | Ado EtOH | -0.65443 | 0.028956 | 4.912e-07 | -0.72528 | -0.58358 |
| Post-hoc Male EPM Time in Open Arms (Fig 2K) | Normal, Identity | Ado EtOH | 0.43031 | 0.3931 | 0.30553 | -0.47619 | 1.3368 |
| EPM Open Arm Entries (Fig 2L) | Normal, Identity | Sex (Male) | -106.73 | 45.694 | 0.034904 | -204.73 | -8.7215 |
|  |  | Ado EtOH | -0.51325 | 0.28469 | 0.092978 | -1.1238 | 0.097356 |
|  |  | Sex x Ado EtOH | 1.0538 | 0.42196 | 0.025593 | 0.14881 | 1.9588 |
| Post-hoc Female EPM Open Arm Entries (Fig 2L) | Normal, Identity | Ado EtOH | -0.55347 | 0.19474 | 0.029487 | -1.03 | -0.076954 |
| Post-hoc Male  EPM Open Arm Entries (Fig 2L) | Normal, Identity | Ado EtOH | 0.54058 | 0.36688 | 0.17885 | -0.30544 | 1.3866 |
| EPM Head Dips (Fig 2O) | Normal, Identity | Sex (Male) | -232.09 | 70.78 | 0.0054862 | -383.9 | -80.283 |
|  |  | Ado EtOH | -0.95715 | 0.44098 | 0.047661 | -1.903 | -0.011331 |
|  |  | Sex x Ado EtOH | 2.1466 | 0.65361 | 0.0054301 | 0.74475 | 3.5485 |
| Post-hoc Female EPM Head Dips (Fig 2O) | Normal, Identity | Ado EtOH | -1.0553 | 0.23463 | 0.0041145 | -1.6294 | -0.48116 |
| Post-hoc Male EPM Head Dips (Fig 2O) | Normal, Identity | Ado EtOH | 1.1895 | 0.56323 | 0.067676 | -0.10934 | 2.4883 |
| EPM Distance Traveled (Fig 2P) | Normal, Identity | Sex (Male) | -1527.4 | 733.9 | 0.056256 | -3101.5 | 46.684 |
|  |  | Ado EtOH | -4.1053 | 4.5725 | 0.38447 | -13.912 | 5.7018 |
|  |  | Sex x Ado EtOH | 13.567 | 6.7773 | 0.065063 | -0.96843 | 28.103 |

**Supp Table 5**: All GLME outcomes for experiment 2 examining adolescent DID effects on adult exploratory behavior (presented in main text Figure 2).

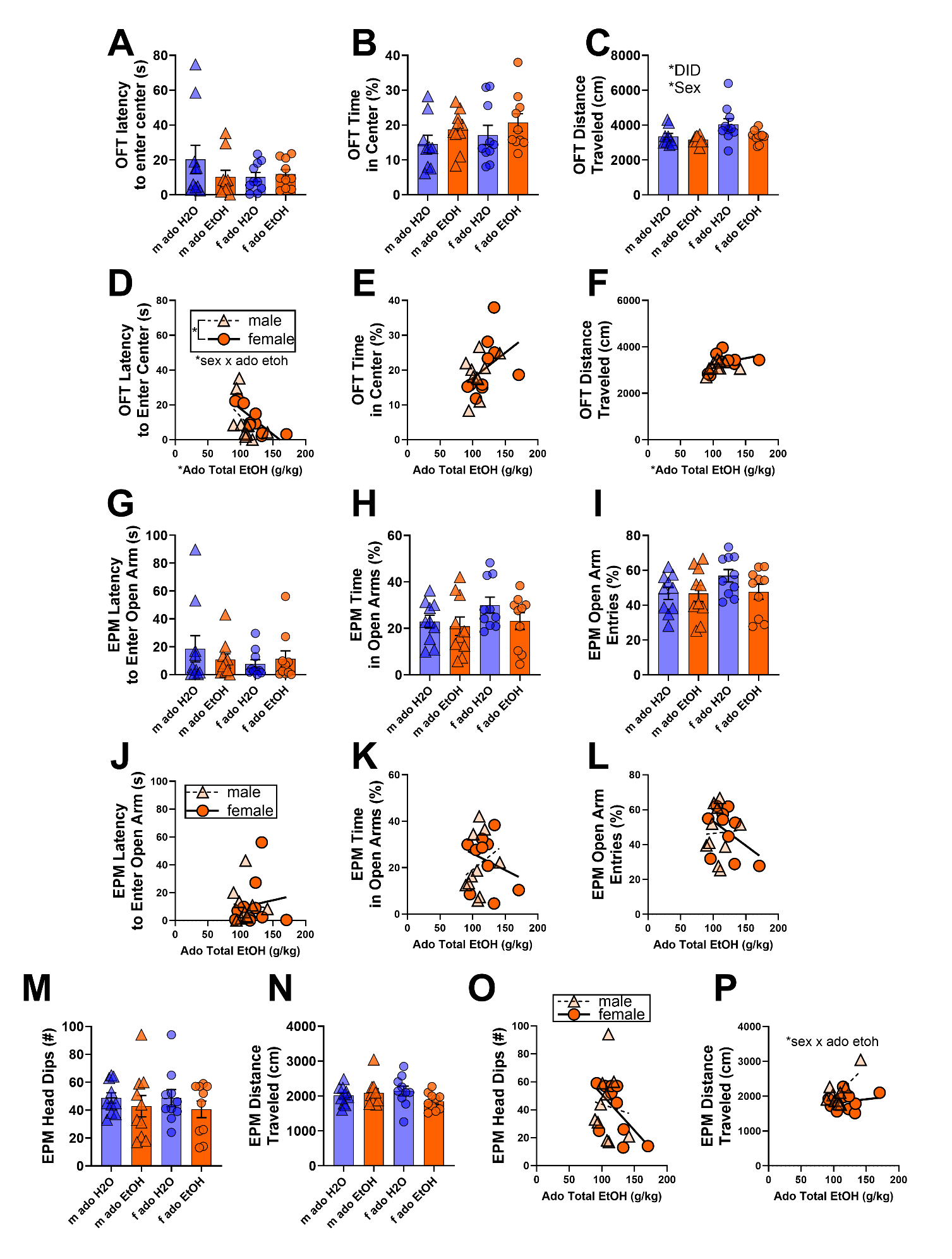
**Supp Fig 3.** OFT and EPM **24 hrs** after **adolescent** DID. Across DID and sex groups, there were no differences detected in **(A)** OFT latency to enter center of arena or **(B)** OFT percent time spent in center of arena. **(C)** OFT total distance traveled was significantly influenced by DID (F_1,35_ = 4.9, *p* = 0.0321) and sex (F_1,35_ = 5.0, *p* = 0.0341). Secondary analyses in only alcohol-exposed subjects found that **(D)** OFT latency to enter center was significantly lower in males than in females (male: -4.5008 ± 1.5347, p = 0.0098), decreased by total alcohol consumed (ado EtOH: -0.03576 ± 0.0082, p = 0.0005), and influenced by a sex by total alcohol interaction (male x ado EtOH: 0.03393 ± 0.01395, p = 0.0271). Post hoc follow-up analyses within sex groups revealed that total alcohol consumption significantly decreased OFT latency to enter the center in females (ado EtOH: -0.2946 ± 0.0658, p = 0.0021) but not in males. **(E)** OFT percent time in center was not associated with adolescent EtOH consumption. **(F)** OFT total distance traveled was significantly increased by total adolescent EtOH consumption (ado EtOH: 8.5896 ± 3.693, p = 0.0335). Across DID and sex groups, there were no differences in **G)** EPM latency to enter an open arm, **(H)** EPM percent time spent in open arms, or **(I)** EPM percent open arm entries. In analyses examining only alcohol-treated subjects, **(J)** EPM latency to enter an open arm, **(K)** EPM percent time spent in open arms, and **(L)** EPM percent open arm entries were not significantly associated with total adolescent EtOH consumption. DID and sex did not influence **M)** EPM number of head dips or **(N)** EPM total distance traveled. In analyses of only alcohol-exposed subjects, **(O)** EPM number of head dips was not associated with total adolescent EtOH consumption. However, **(P)** EPM total distance traveled was influenced by an interaction of total adolescent EtOH consumption and sex (male x ado EtOH: 16.718 ± 6.5302, p = 0.0210). Post hoc follow-up analyses within sex groups revealed that EPM total distance traveled was significantly increased by total adolescent alcohol consumption in males (ado EtOH: 18.88 ± 5.726, p = 0.0110) but not in females. M = male; f = female; ado = adolescent. In scatter plots, a simple linear regression was used to generate lines of best fit and aid interpretation but were independent from statistical analysis. Data are shown as mean +/- SEM. *p<0.05. n=9-10/sex/treatment.

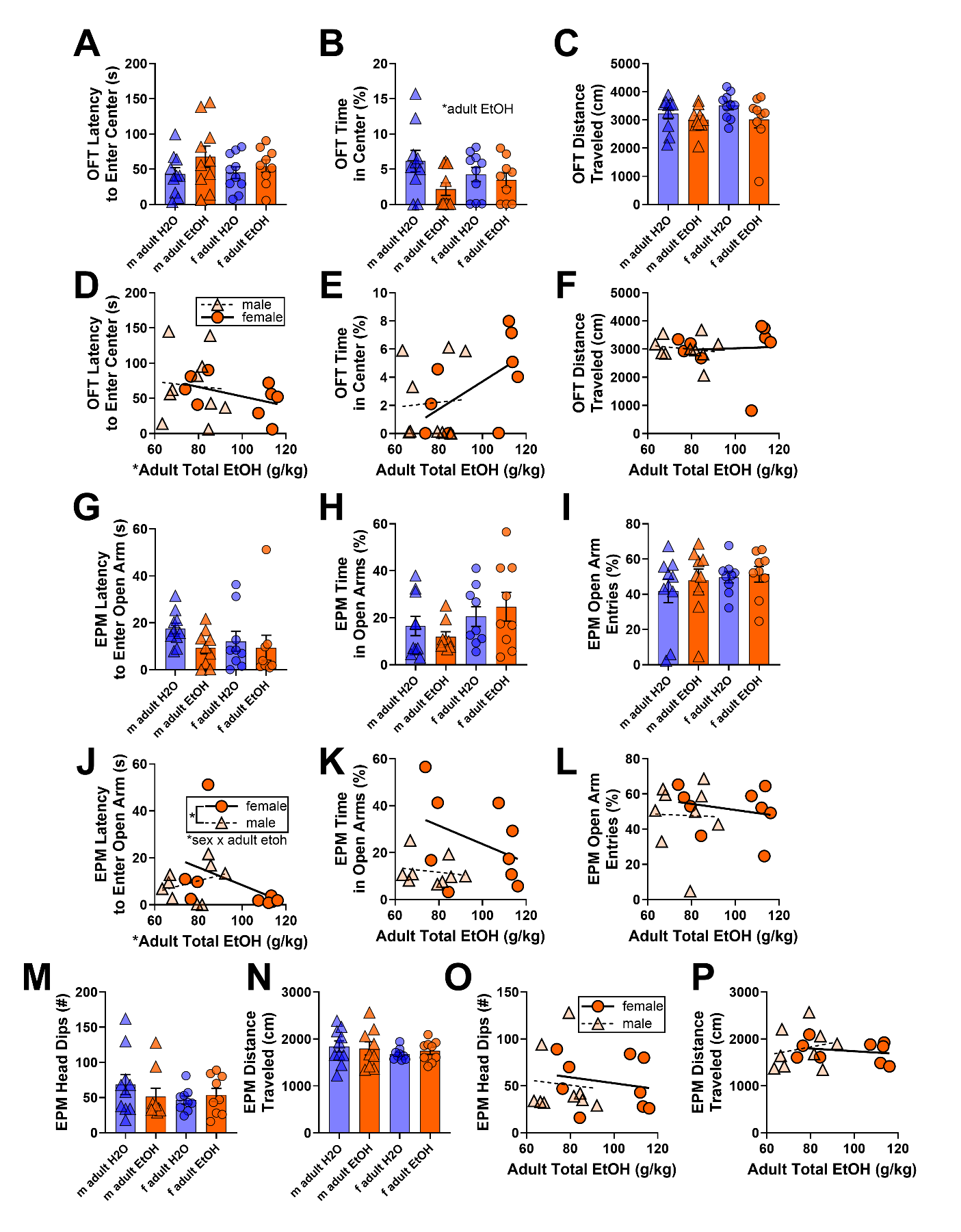
**Supp Fig 4.** OFT and EPM **30 days** after **adult** DID. Across DID and sex groups, there were no differences found for **A)** OFT latency to enter center of arena. However, **(B)** OFT percent time spent in center of arena was significantly lowered after adult DID (F_1,35_ = 4.3, *p* = 0.0450). **(C)** DID and sex did not influence OFT total distance traveled. In separate analyses testing effects of total alcohol consumed among only alcohol-exposed mice, **(D)** OFT latency to enter center was decreased by total alcohol consumed (adult EtOH: -0.0122 ± 0.0026, p = 0.0003). Total alcohol consumed did not significantly alter **(E)** OFT percent time in center or **(F)** OFT total distance traveled compared to adult EtOH consumption. DID and sex did not influence **G)** EPM latency to enter an open arm, **(H)** EPM percent time spent in open arms, or **(I)** EPM percent open arm entries. Analyses within alcohol-treated groups revealed that **(J)** EPM latency to enter an open arm was significantly lower in males than in females (male: -3.2651 ± 1.1546, p = 0.0134), decreased by total alcohol consumed (adult EtOH: -0.0339 ± 0.0081, p = 0.0009), and influenced by a sex by total alcohol interaction (male x adult EtOH: 0.0370 ± 0.0140, p = 0.0193). Follow-up post hoc comparisons within sex groups found that total adult alcohol consumption significantly decreased EPM latency to enter an open arm in females (adult EtOH: -0.0354 ± 0.0082, p = 0.0035) but not in males. Total adult alcohol consumption was not predictive of **(K)** EPM percent time spent in open arms and **(L)** EPM percent open arm entries. Across DID and sex groups, differences were not found in **M)** EPM number of head dips or **(N)** EPM total distance traveled. Total adult alcohol consumption was not associated with **(O)** EPM number of head dips or **(P)** EPM total distance traveled. M = male; f = female. In scatter plots, a simple linear regression was used to generate lines of best fit and aid interpretation but were independent from statistical analysis. Data are shown as mean +/- SEM. *p<0.05. n=9-10/sex/treatment.

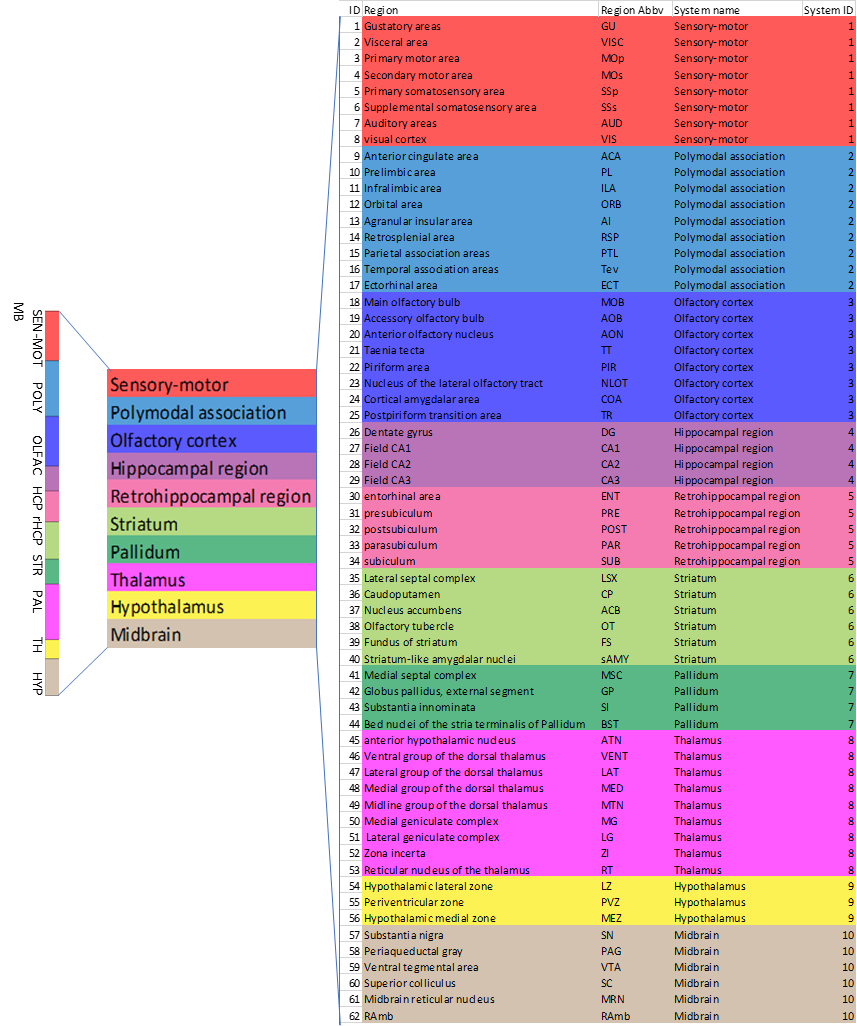

**Supp Fig 5.** Regions of interest (ROIs) and their associated brain systems were defined based on anatomical classifications from the Allen Mouse Brain Atlas (Lein et al., 2007).

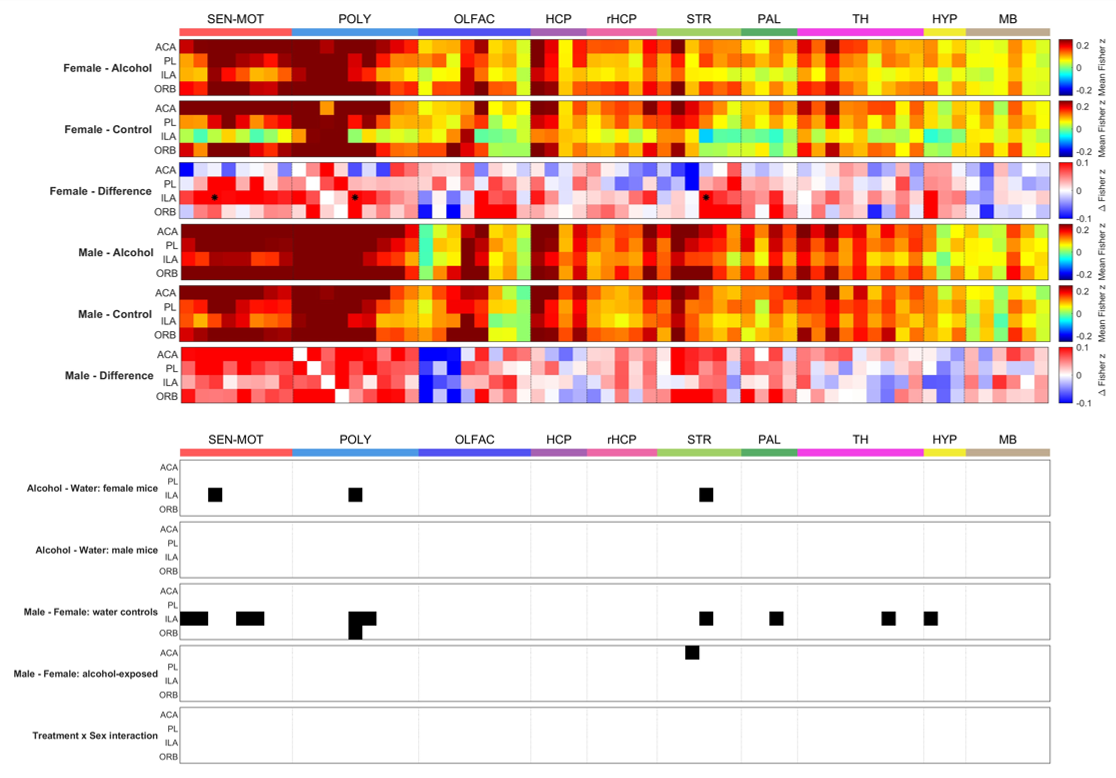

**Supp Fig 6.** Using linear mixed modeling and a significance threshold of p<0.005, pairwise comparisons of RSFC were performed for each ROI within the medial prefrontal cortex. ACA: anterior cingulate area, PL: prelimbic area, ILA: infralimbic area, ORB: orbital area. n=11 mice/sex/treatment.

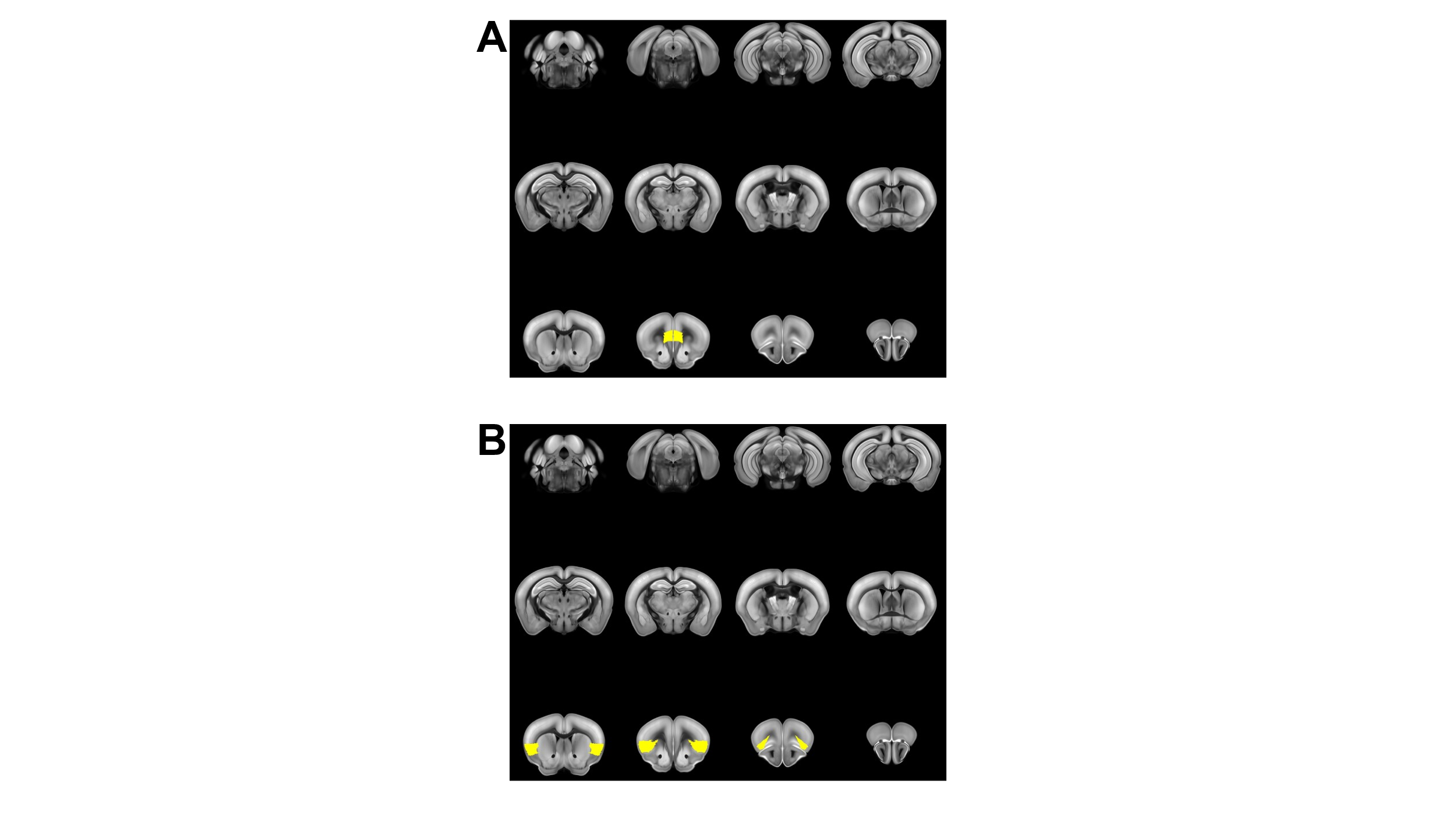
**Supp Fig 7.** Infralimbic cortex (A) and anterior insular cortex (B) seed regions are highlighted in yellow. Seed regions in the mouse brain were defined based on anatomical classifications from the Allen Mouse Brain Atlas (Lein et al., 2007).

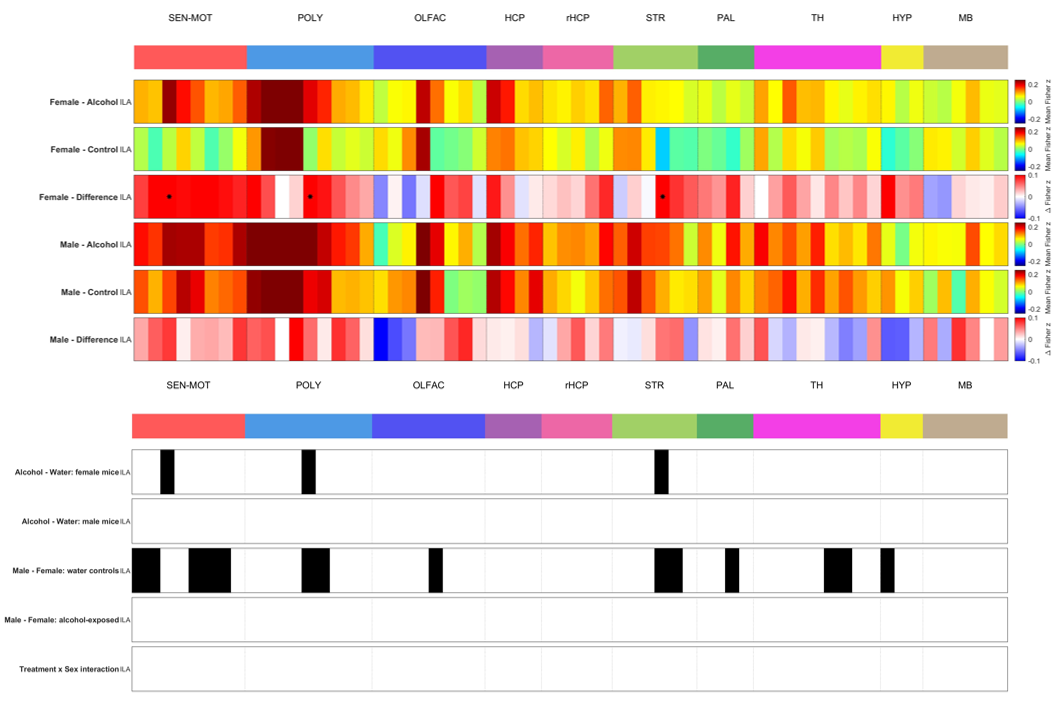

**Supp Fig 8.** Using linear mixed modeling with an FDR correction and a significance threshold of q<0.05, pairwise comparisons of RSFC were performed for the infralimbic area with each ROI. This analysis extends that shown in Supp Fig 6 by applying FDR correction for the infralimbic area with each ROI. Significant effects of adolescent alcohol exposure were found only in females, and basal sex differences independent of alcohol exposure were also identified. Notably, significant effects of adolescent alcohol exposure were found in female mice for RSFC between the infralimbic cortex and the primary motor area, agranular insular area, and olfactory tubercle. n=11 mice/sex/treatment.

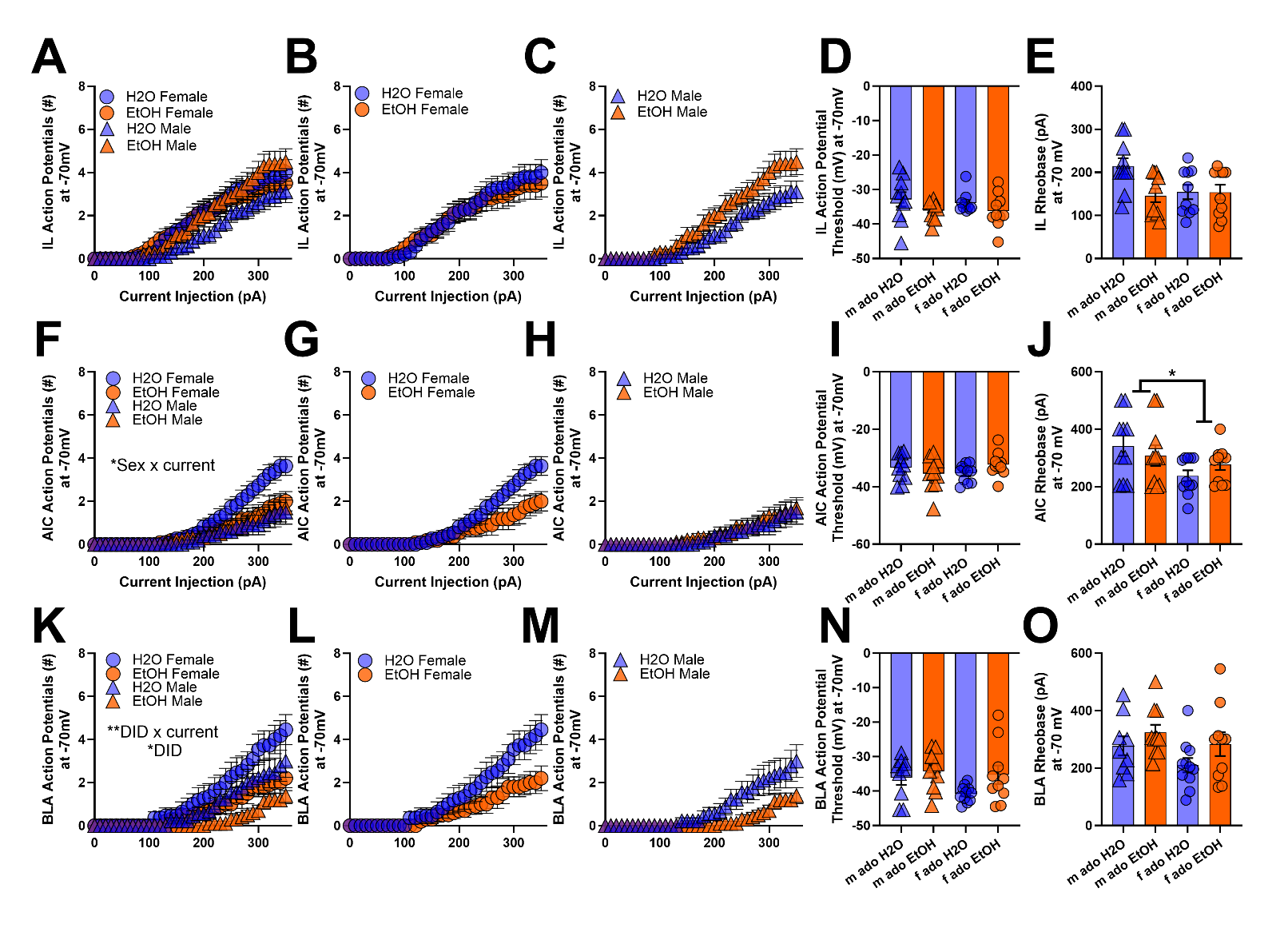
**Supp Fig 9. Adolescent alcohol consumption has subtle effects on pyramidal cell excitability when cells are held at a common voltage (-70mV).** **(A-C)** Current-induced firing of infralimbic cells at -70mV, with sexes visualized together **(A)** and separately **(B-C)**. **(D-E)** Action potential threshold and rheobase of infralimbic cells at -70mV. **(F-H)** Current-induced firing of anterior insular cortex cells at -70mV, with sexes visualized together **(F)** and separately **(G-H)**. An interaction of sex and current injection was detected for current-induced firing (F_2, 58_ = 4.0, p = 0.0327)**. (I-J)** Action potential threshold and rheobase of anterior insular cortex cells at -70mV. Rheobase was higher in males than in females (F_1, 38_ = 5.8, p = 0.0213). **(K-M)** Current-induced firing of basolateral amygdala cells at -70mV, with sexes visualized together **(K)** and separately **(L-M)**. Adolescent alcohol exposure significantly reduced action potential firing (F_1, 36_ = 5.9, *p* = 0.0203) and in a current-dependent manner (F_2, 58_ =6.4, p = 0.0055). **(N-O)** Action potential threshold and rheobase of basolateral amygdala cells at -70mV. As expected, across all regions, current-induced firing was significantly influenced by current injection (infralimbic: F_2, 64_ = 124.2, *p* < 0.0001; anterior insular cortex: F_2, 58_ = 47.1, *p* < 0.0001; basolateral amygdala: F_2, 58_ = 47.9, *p* < 0.0001).M = male; f = female; ado = adolescent. Data are shown as mean +/- SEM. *p<0.05. **p<0.01. n=6-9 mice/sex/treatment for electrophysiology.

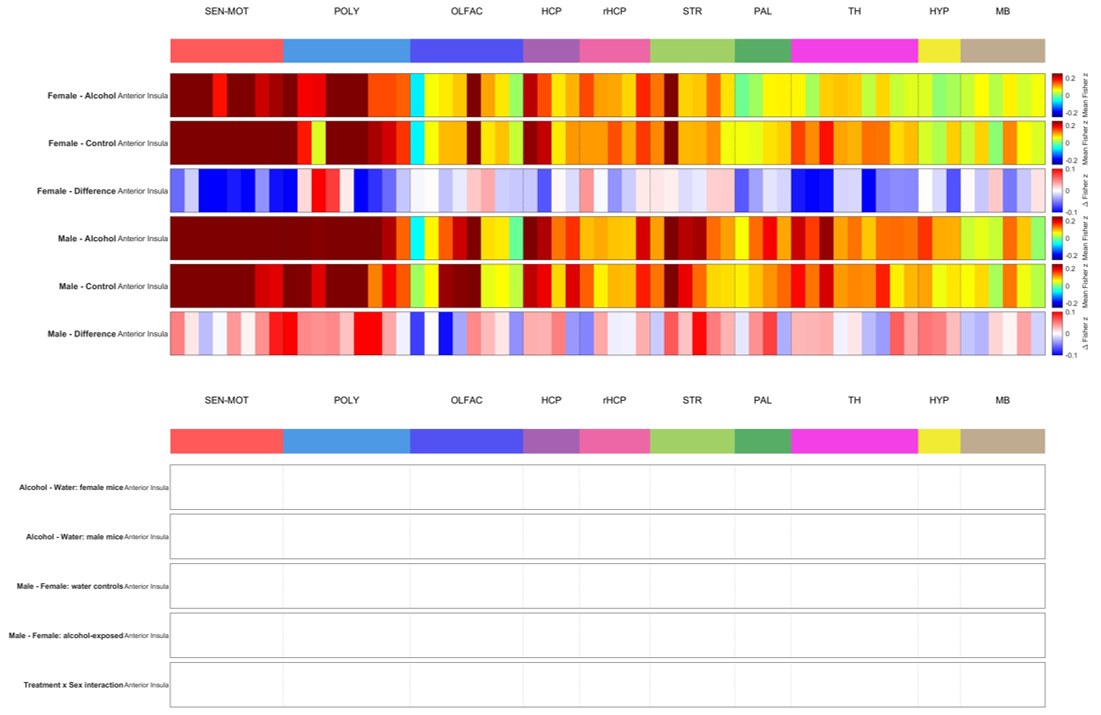

**Supp Fig 10.** Using linear mixed modeling with an FDR correction and a significance threshold of q<0.05, pairwise comparisons of RSFC were performed for the anterior insula with each ROI. No significant treatment or sex effects were found. n=11 mice/sex/treatment.

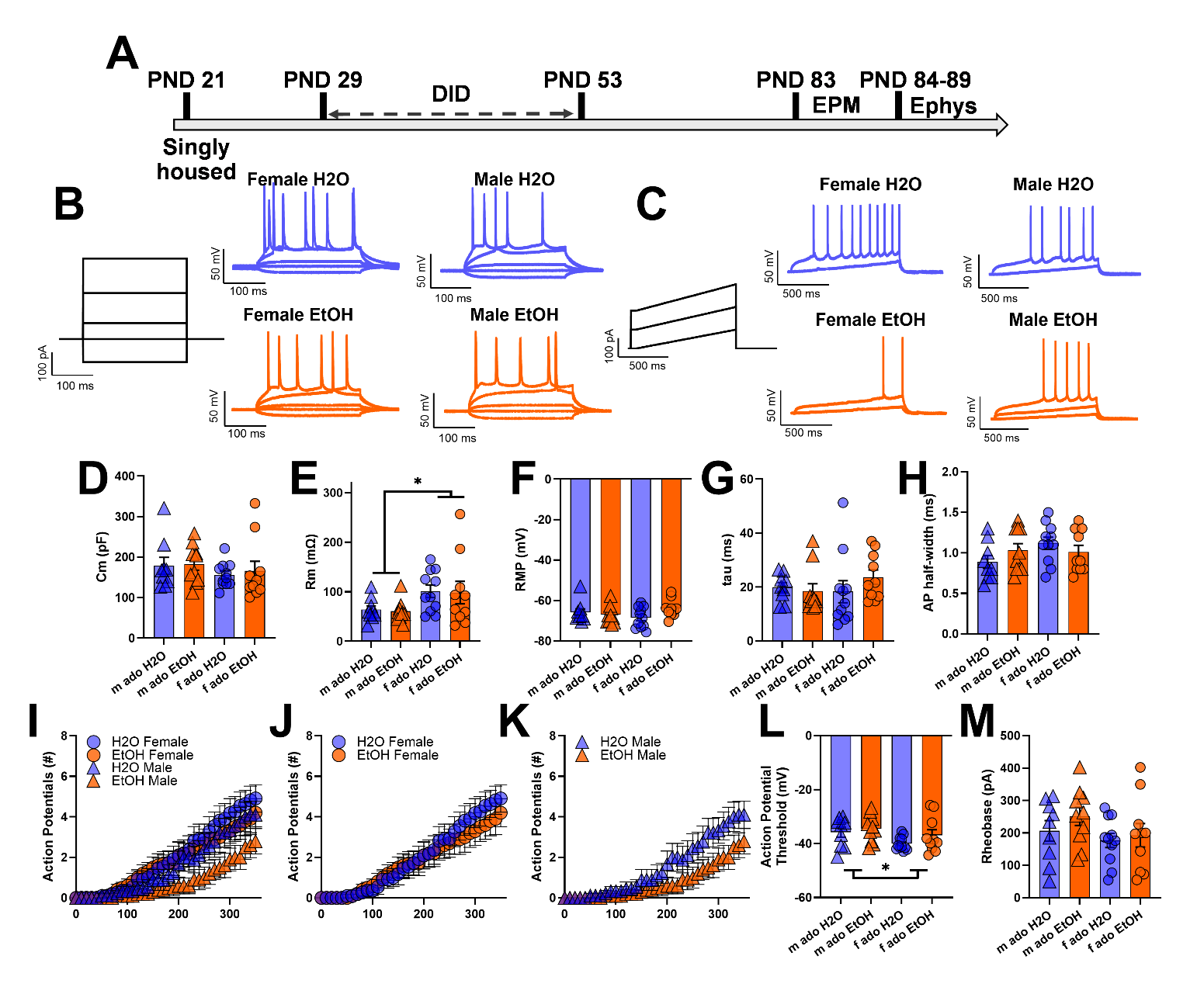

**Supp Fig 11. Basolateral amygdala pyramidal cell excitability after adolescent alcohol consumption.** **(A)** Experimental timeline for patch clamp electrophysiology experiments testing the intrinsic excitability of basolateral amygdala pyramidal cells. **(B)** Representative traces from current-induced firing experiments. **(C)** Representative traces from rheobase recordings. **(D-H)** Cellular capacitance, membrane resistance, resting membrane potential (RMP), time constant (tau), and action potential (AP) half-width**.** Membrane resistance was significantly higher in females than males, independent of alcohol exposure (F_1,36_ = 6.9, *p* = 0.0123). **(I-K)** Current-induced firing of cells at RMP, with sexes visualized together **(I)** and separately **(J-K)**. As expected, current had a significant impact on firing (F_2, 64_ = 69.0, *p* < 0.0001). **(L-M)** Action potential threshold and rheobase of cells at RMP. The action potential threshold was significantly lower in females than in males (F_1,36_ = 4.4, *p* = 0.0431). M = male; f = female; ado = adolescent. Data are shown as mean +/- SEM. *p<0.05. **p<0.01. n=11 mice/sex/treatment for fMRI and n=7-9 mice/sex/treatment for electrophysiology.

| **Dependent variable** | **Effect** | **F (DFn, DFd)** | **p** |
| --- | --- | --- | --- |
| IL Cm – (Fig 4F)  2-way (DID, Sex) ANOVA | DID | F (1, 36) = 5.760 | P=0.0217 |
|  | Sex | F (1, 36) = 0.05959 | P=0.8085 |
|  | DID x Sex | F (1, 36) = 0.8299 | P=0.3683 |
| IL Rm – (Fig 4G)  2-way (DID, Sex) ANOVA | DID | F (1, 36) = 3.841 | P=0.0578 |
|  | Sex | F (1, 36) = 1.911 | P=0.1754 |
|  | DID x Sex | F (1, 36) = 1.004 | P=0.3230 |
| IL RMP – (Fig 4H)  2-way (DID, Sex) ANOVA | DID | F (1, 36) = 2.830 | P=0.1012 |
|  | Sex | F (1, 36) = 0.02245 | P=0.8817 |
|  | DID x Sex | F (1, 36) = 8.939 | P=0.0050 |
| IL tau – (Fig 4I)  2-way (DID, Sex) ANOVA | DID | F (1, 36) = 0.2894 | P=0.5939 |
|  | Sex | F (1, 36) = 1.110 | P=0.2990 |
|  | DID x Sex | F (1, 36) = 0.3826 | P=0.5401 |
| IL AP half width – (Fig 4J)  2-way (DID, Sex) ANOVA | DID | F (1, 36) = 0.002477 | P=0.9606 |
|  | Sex | F (1, 36) = 5.948 | P=0.0198 |
|  | DID x Sex | F (1, 36) = 0.5574 | P=0.4602 |
| IL current-induced firing – (Fig 4K)  3-way (DID, Sex, Current) ANOVA | DID | F (1, 36) = 3.322 | P=0.0767 |
|  | Sex | F (1, 36) = 0.4966 | P=0.4855 |
|  | Current | F (2.063, 74.29) = 181.4 | P<0.0001 |
|  | DID x Current | F (2.063, 74.29) = 1.272 | P=0.2869 |
|  | Sex x Current | F (2.063, 74.29) = 0.6071 | P=0.5526 |
|  | DID x Sex | F (1, 36) = 2.906 | P=0.0969 |
|  | DID x Sex x Current | F (2.063, 74.29) = 6.636 | P=0.0020 |
| IL current-induced firing  Post hoc – Females (Fig 4L)  2-way (DID, Current) ANOVA | DID | F (1, 18) = 0.005719 | P=0.9406 |
|  | Current | F (2.083, 37.49) = 93.25 | P<0.0001 |
|  | DID x Current | F (2.083, 37.49) = 2.291 | P=0.1132 |
| IL current-induced firing  Post hoc – Males (Fig 4M)  2-way (DID, Current) ANOVA | DID | F (1, 18) = 7.971 | P=0.0113 |
|  | Current | F (1.897, 34.15) = 89.06 | P<0.0001 |
|  | DID x Current | F (1.897, 34.15) = 5.423 | P=0.0100 |
| IL AP threshold – (Fig 4N)  2-way (DID, Sex) ANOVA | DID | F (1, 36) = 0.3143 | P=0.5785 |
|  | Sex | F (1, 36) = 0.3639 | P=0.5501 |
|  | DID x Sex | F (1, 36) = 5.037e-005 | P=0.9944 |
| IL rheobase – (Fig 4O)  2-way (DID, Sex) ANOVA | DID | F (1, 36) = 3.616 | P=0.0652 |
|  | Sex | F (1, 36) = 2.633 | P=0.1134 |
|  | DID x Sex | F (1, 36) = 0.003587 | P=0.9526 |
| IL AP threshold at -70 mV – (Supp Fig 9)  2-way (DID, Sex) ANOVA | DID | F (1, 36) = 3.616 | P=0.0652 |
|  | Sex | F (1, 36) = 0.1554 | P=0.6958 |
|  | DID x Sex | F (1, 36) = 0.1056 | P=0.7471 |
| IL rheobase at -70 mV – (Supp Fig 9)  2-way (DID, Sex) ANOVA | DID | F (1, 36) = 4.085 | P=0.0507 |
|  | Sex | F (1, 36) = 2.414 | P=0.1290 |
|  | DID x Sex | F (1, 36) = 4.021 | P=0.0525 |
| IL current-induced firing at -70mV –  (Supp Fig 9)  3-way (DID, Sex, Current) ANOVA | DID | F (1, 36) = 1.083 | P=0.3050 |
|  | Sex | F (1, 36) = 0.8016 | P=0.3766 |
|  | Current | F (1.771, 63.77) = 124.2 | P<0.0001 |
|  | DID x Current | F (1.771, 63.77) = 0.6629 | P=0.5013 |
|  | Sex x Current | F (1.771, 63.77) = 1.199 | P=0.3041 |
|  | DID x Sex | F (1, 36) = 1.720 | P=0.1980 |
|  | DID x Sex x Current | F (1.771, 63.77) = 2.146 | P=0.1309 |

**Supp Table 6**: All ANOVA outcomes for the analysis of infralimbic excitability after adolescent DID (presented in main text Figure 4).

| **Dependent variable** | **Effect** | **F (DFn, DFd)** | **p** |
| --- | --- | --- | --- |
| AIC Cm – (Fig 5F)  2-way (DID, Sex) ANOVA | DID | F (1, 38) = 3.067 | P=0.0880 |
|  | Sex | F (1, 38) = 1.983 | P=0.1672 |
|  | DID x Sex | F (1, 38) = 0.03085 | P=0.8615 |
| AIC Rm – (Fig 5G)  2-way (DID, Sex) ANOVA | DID | F (1, 38) = 0.02994 | P=0.8635 |
|  | Sex | F (1, 38) = 0.06719 | P=0.7969 |
|  | DID x Sex | F (1, 38) = 2.818 | P=0.1014 |
| AIC RMP – (Fig 5H)  2-way (DID, Sex) ANOVA | DID | F (1, 38) = 0.01857 | P=0.8923 |
|  | Sex | F (1, 38) = 0.01072 | P=0.9181 |
|  | DID x Sex | F (1, 38) = 0.4529 | P=0.5050 |
| AIC tau – (Fig 5I)  2-way (DID, Sex) ANOVA | DID | F (1, 38) = 1.114 | P=0.2978 |
|  | Sex | F (1, 38) = 1.271 | P=0.2666 |
|  | DID x Sex | F (1, 38) = 0.9306 | P=0.3408 |
| AIC AP half width – (Fig 5J)  2-way (DID, Sex) ANOVA | DID | F (1, 38) = 6.883 | P=0.0125 |
|  | Sex | F (1, 38) = 4.235 | P=0.0465 |
|  | DID x Sex | F (1, 38) = 0.04171 | P=0.8393 |
| AIC current-induced firing – (Fig 5K)  3-way (DID, Sex, Current) ANOVA | DID | F (1, 38) = 0.1234 | P=0.7274 |
|  | Sex | F (1, 38) = 2.890 | P=0.0973 |
|  | Current | F (1.762, 66.95) = 88.42 | P<0.0001 |
|  | DID x Current | F (1.762, 66.95) = 0.5712 | P=0.5463 |
|  | Sex x Current | F (1.762, 66.95) = 4.713 | P=0.0154 |
|  | DID x Sex | F (1, 38) = 0.5921 | P=0.4463 |
|  | DID x Sex x Current | F (1.762, 66.95) = 0.3566 | P=0.6743 |
| AIC AP threshold – (Fig 5N)  2-way (DID, Sex) ANOVA | DID | F (1, 38) = 0.6732 | P=0.4170 |
|  | Sex | F (1, 38) = 0.1622 | P=0.6894 |
|  | DID x Sex | F (1, 38) = 0.6053 | P=0.4414 |
| AIC rheobase – (Fig 5O)  2-way (DID, Sex) ANOVA | DID | F (1, 38) = 0.2584 | P=0.6142 |
|  | Sex | F (1, 38) = 2.983 | P=0.0922 |
|  | DID x Sex | F (1, 38) = 1.177 | P=0.2848 |
| AIC AP threshold at -70 mV – (Supp Fig 9)  2-way (DID, Sex) ANOVA | DID | F (1, 38) = 0.1076 | P=0.7447 |
|  | Sex | F (1, 38) = 0.1750 | P=0.6780 |
|  | DID x Sex | F (1, 38) = 3.522 | P=0.0682 |
| AIC rheobase at -70 mV – (Supp Fig 9)  2-way (DID, Sex) ANOVA | DID | F (1, 38) = 0.009461 | P=0.9230 |
|  | Sex | F (1, 38) = 5.774 | P=0.0213 |
|  | DID x Sex | F (1, 38) = 1.657 | P=0.2057 |
| AIC current-induced firing at -70mV –  (Supp Fig 9)  3-way (DID, Sex, Current) ANOVA | DID | F (1, 38) = 0.9438 | P=0.3374 |
|  | Sex | F (1, 38) = 3.575 | P=0.0663 |
|  | Current | F (1.538, 58.43) = 47.91 | P<0.0001 |
|  | DID x Current | F (1.538, 58.43) = 1.864 | P=0.1722 |
|  | Sex x Current | F (1.538, 58.43) = 4.023 | P=0.0327 |
|  | DID x Sex | F (1, 38) = 1.909 | P=0.1751 |
|  | DID x Sex x Current | F (1.538, 58.43) = 2.484 | P=0.1050 |

**Supp Table 7**: All ANOVA outcomes for the analysis of anterior insular cortex excitability after adolescent DID (presented in main text Figure 5).

| **Dependent variable** | **Effect** | **F (DFn, DFd)** | **p** |
| --- | --- | --- | --- |
| BLA Cm – (Supp Fig 10D)  2-way (DID, Sex) ANOVA | DID | F (1, 36) = 0.1501 | P=0.7007 |
|  | Sex | F (1, 36) = 1.287 | P=0.2641 |
|  | DID x Sex | F (1, 36) = 0.02676 | P=0.8710 |
| BLA Rm – (Supp Fig 10E)  2-way (DID, Sex) ANOVA | DID | F (1, 36) = 0.05518 | P=0.8156 |
|  | Sex | F (1, 36) = 6.941 | P=0.0123 |
|  | DID x Sex | F (1, 36) = 0.0005702 | P=0.9811 |
| BLA RMP – (Supp Fig 10F)  2-way (DID, Sex) ANOVA | DID | F (1, 36) = 1.143 | P=0.2921 |
|  | Sex | F (1, 36) = 0.02186 | P=0.8833 |
|  | DID x Sex | F (1, 36) = 3.505 | P=0.0693 |
| BLA tau – (Supp Fig 10G)  2-way (DID, Sex) ANOVA | DID | F (1, 36) = 0.4099 | P=0.5261 |
|  | Sex | F (1, 36) = 0.3263 | P=0.5714 |
|  | DID x Sex | F (1, 36) = 1.248 | P=0.2713 |
| BLA AP half width – (Supp Fig 10H)  2-way (DID, Sex) ANOVA | DID | F (1, 36) = 0.04230 | P=0.8382 |
|  | Sex | F (1, 36) = 1.709 | P=0.1994 |
|  | DID x Sex | F (1, 36) = 2.425 | P=0.1282 |
| BLA current-induced firing – (Supp Fig 10I)  3-way (DID, Sex, Current) ANOVA | DID | F (1, 36) = 1.442 | P=0.2377 |
|  | Sex | F (1, 36) = 3.125 | P=0.0856 |
|  | Current | F (1.764, 63.51) = 68.96 | P<0.0001 |
|  | DID x Current | F (1.764, 63.51) = 2.415 | P=0.1039 |
|  | Sex x Current | F (1.764, 63.51) = 2.100 | P=0.1364 |
|  | DID x Sex | F (1, 36) = 0.5246 | P=0.4736 |
|  | DID x Sex x Current | F (1.764, 63.51) = 0.3737 | P=0.6633 |
| BLA AP threshold – (Supp Fig 10L)  2-way (DID, Sex) ANOVA | DID | F (1, 36) = 1.884 | P=0.1783 |
|  | Sex | F (1, 36) = 4.398 | P=0.0431 |
|  | DID x Sex | F (1, 36) = 0.3041 | P=0.5847 |
| BLA rheobase – (Supp Fig 10M)  2-way (DID, Sex) ANOVA | DID | F (1, 36) = 1.054 | P=0.3113 |
|  | Sex | F (1, 36) = 2.244 | P=0.1429 |
|  | DID x Sex | F (1, 36) = 0.2326 | P=0.6325 |
| BLA AP threshold at -70 mV – (Supp Fig 9)  2-way (DID, Sex) ANOVA | DID | F (1, 36) = 3.676 | P=0.0632 |
|  | Sex | F (1, 36) = 2.270 | P=0.1406 |
|  | DID x Sex | F (1, 36) = 0.9203 | P=0.3438 |
| BLA rheobase at -70 mV – (Supp Fig 9)  2-way (DID, Sex) ANOVA | DID | F (1, 36) = 3.494 | P=0.0697 |
|  | Sex | F (1, 36) = 2.804 | P=0.1027 |
|  | DID x Sex | F (1, 36) = 0.1664 | P=0.6858 |
| BLA current-induced firing – (Supp Fig 9)  3-way (DID, Sex, Current) ANOVA | DID | F (1, 36) = 5.897 | P=0.0203 |
|  | Sex | F (1, 36) = 3.634 | P=0.0646 |
|  | Current | F (1.599, 57.57) = 47.12 | P<0.0001 |
|  | DID x Current | F (1.599, 57.57) = 6.424 | P=0.0055 |
|  | Sex x Current | F (1.599, 57.57) = 2.711 | P=0.0862 |
|  | DID x Sex | F (1, 36) = 0.03342 | P=0.8560 |
|  | DID x Sex x Current | F (1.599, 57.57) = 0.2579 | P=0.7236 |

**Supp Table 8**: All ANOVA outcomes for the analysis of basolateral amygdala excitability after adolescent DID (presented in Supp Fig 11).
